## Supporting Information for "Synthesis of 5-Benzylamino and 5-Alkylamino-substituted Pyrimido[4,5-c]quinoline Derivatives as CSNK2A Inhibitors with Antiviral Activity"

[1] *Structural Genomics Consortium, UNC Eshelman School of Pharmacy, University of North Carolina at Chapel Hill, Chapel Hill, NC 27599, United States;* (K.H.A); (J.L.S); (A.D.A); (T.M.W)
[2] *Rapidly Emerging Antiviral Drug Development Initiative (READDI), Chapel Hill, NC 27599, United States;* (R.J.D); (N.J.M); (A.D.A); (T.M.W).
[3] *Department of Microbiology & Immunology, University of North Carolina at Chapel Hill, Chapel Hill, NC 27599, United States;* (R.J.D); (N.J.M).
[4] *Lineberger Comprehensive Cancer Center, University of North Carolina at Chapel Hill, Chapel Hill, NC 27599, United States.* (R.J.D); (N.J.M).
[5] *Department of Chemistry, University of North Carolina at Chapel Hill, Chapel Hill, NC 27599, United States.* (R.J.D).
[6] *Laboratory for Molecular Modeling, Division of Chemical Biology and Medicinal Chemistry, UNC Eshelman School of Pharmacy, University of North Carolina, Chapel Hill, NC 27599, United States*; (J.W)
[7] *Center of Integrative Chemical Biology and Drug Discovery, UNC Eshelman School of Pharmacy, University of North Carolina, Chapel Hill, NC 27599, United States.* (K.I.P)

**Table of Contents** **Page**

| | |
|---|---|
| Figure S1. $IC_{50}$ curves for CSNK2 NanoBRET assays | S2–3 |
| Figure S2. $IC_{50}$ curves for MHV-NLuc assays | S4 |
| Figures S3-S37 NMR spectra of all final compounds | S5–39 |

**Figure S1.** $IC_{50}$ curves for CSNK2 NanoBRET assays

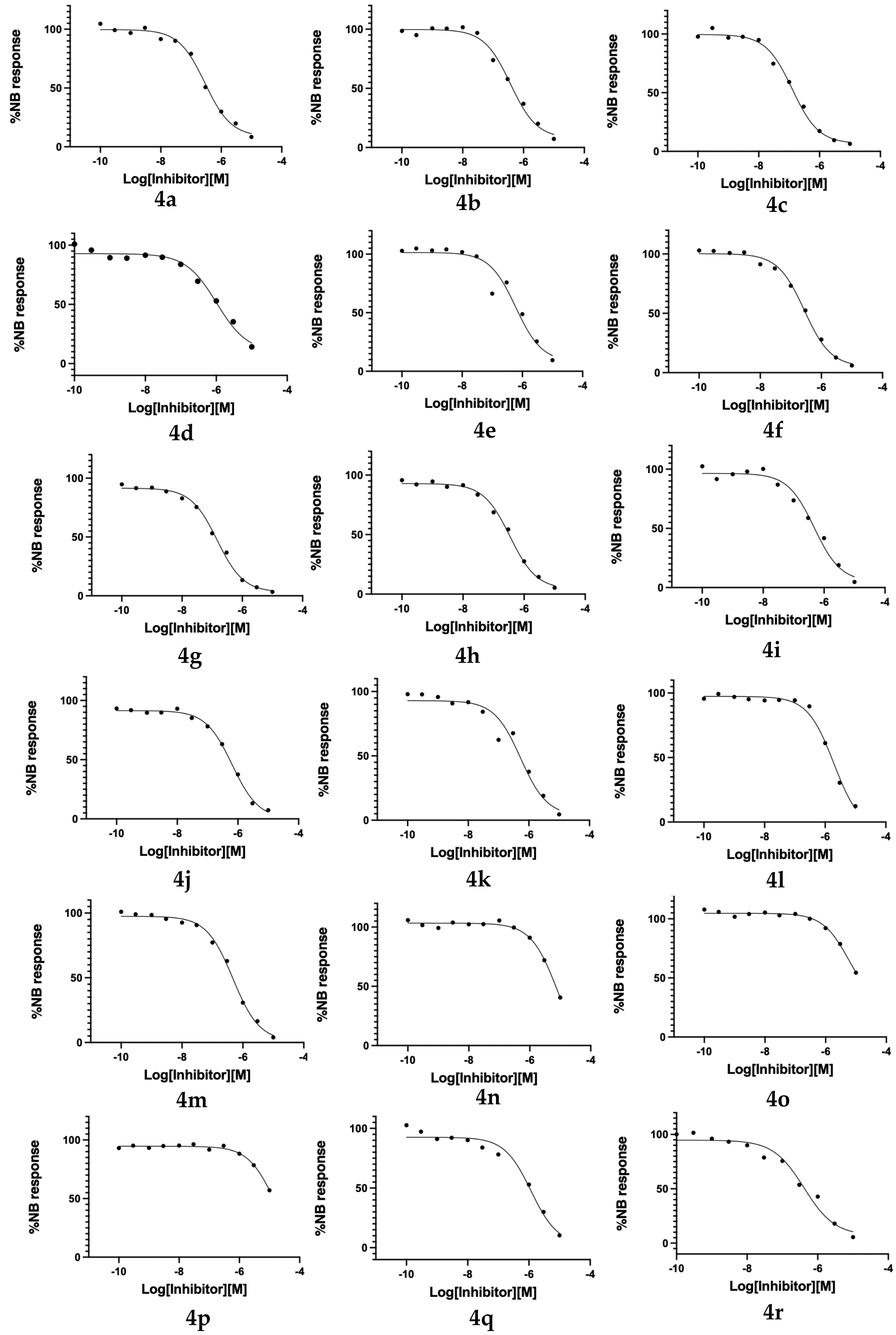

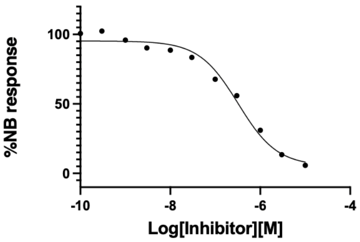


**4s**

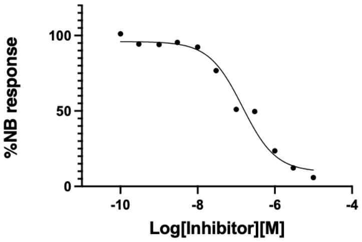


**4t**

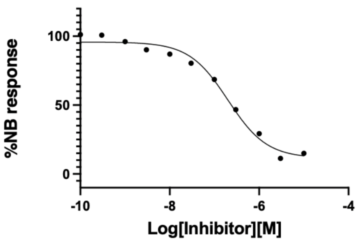


**4u**

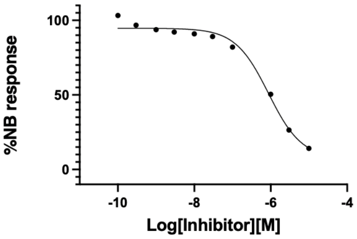


**4v**

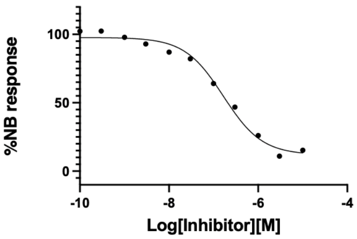


**4w**

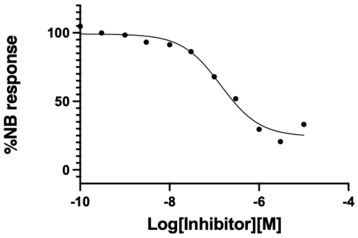


**4x**

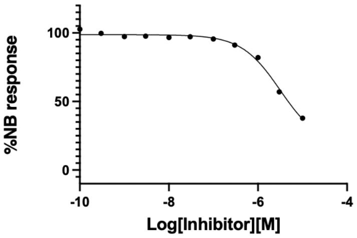


**4y**

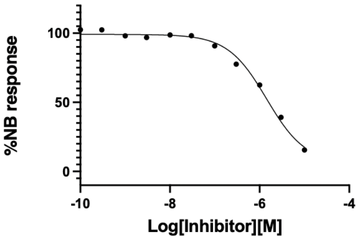


**4z**

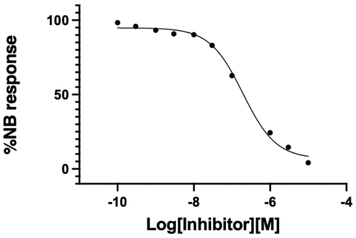


**4aa**

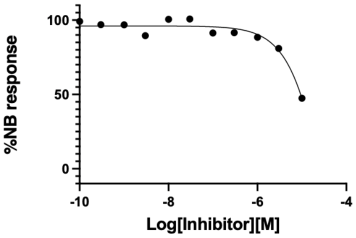


**4ab**

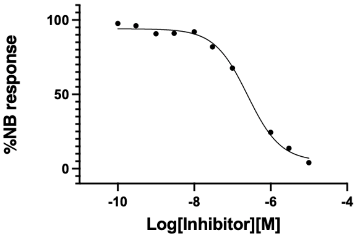


**4ac**

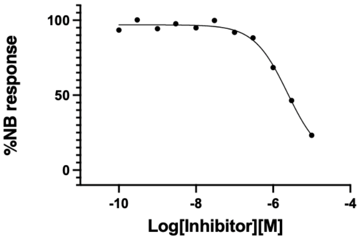


**4ad**

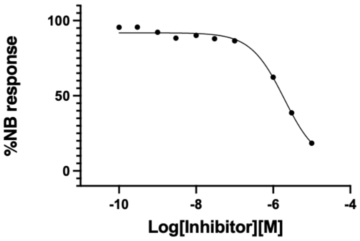


**4ae**

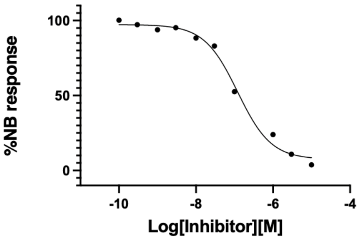


**4af**

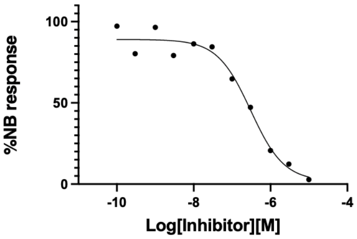


**4ag**

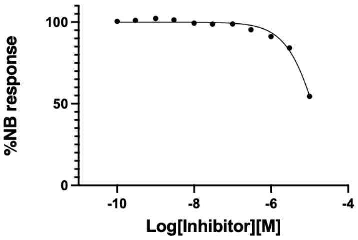


**4ah**

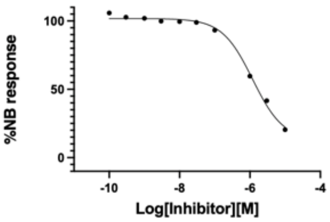


**4ai**

**Figure S2.** $IC_{50}$ curves for MHV-NLuc assays

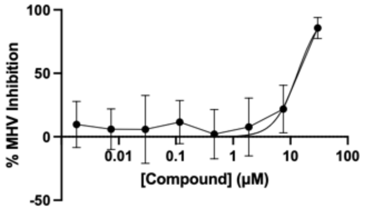


**4c**

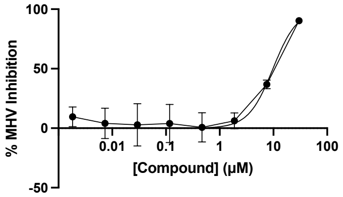


**4g**

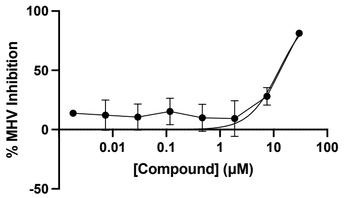


**4t**

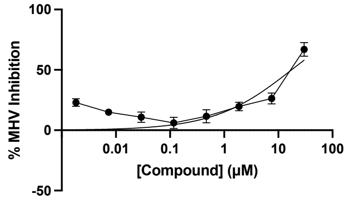


**4u**

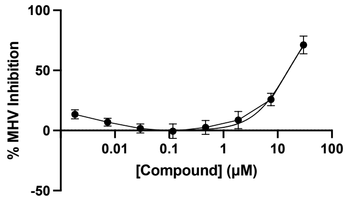


**4w**

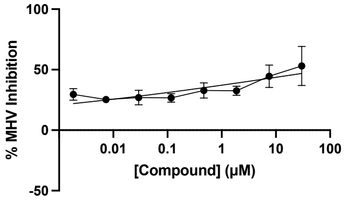


**4x**

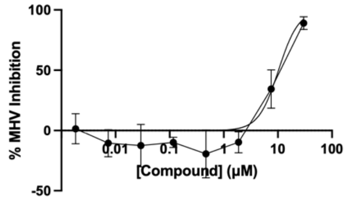


**4aa**

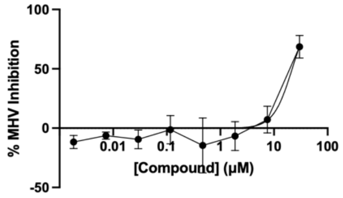


**4ag**

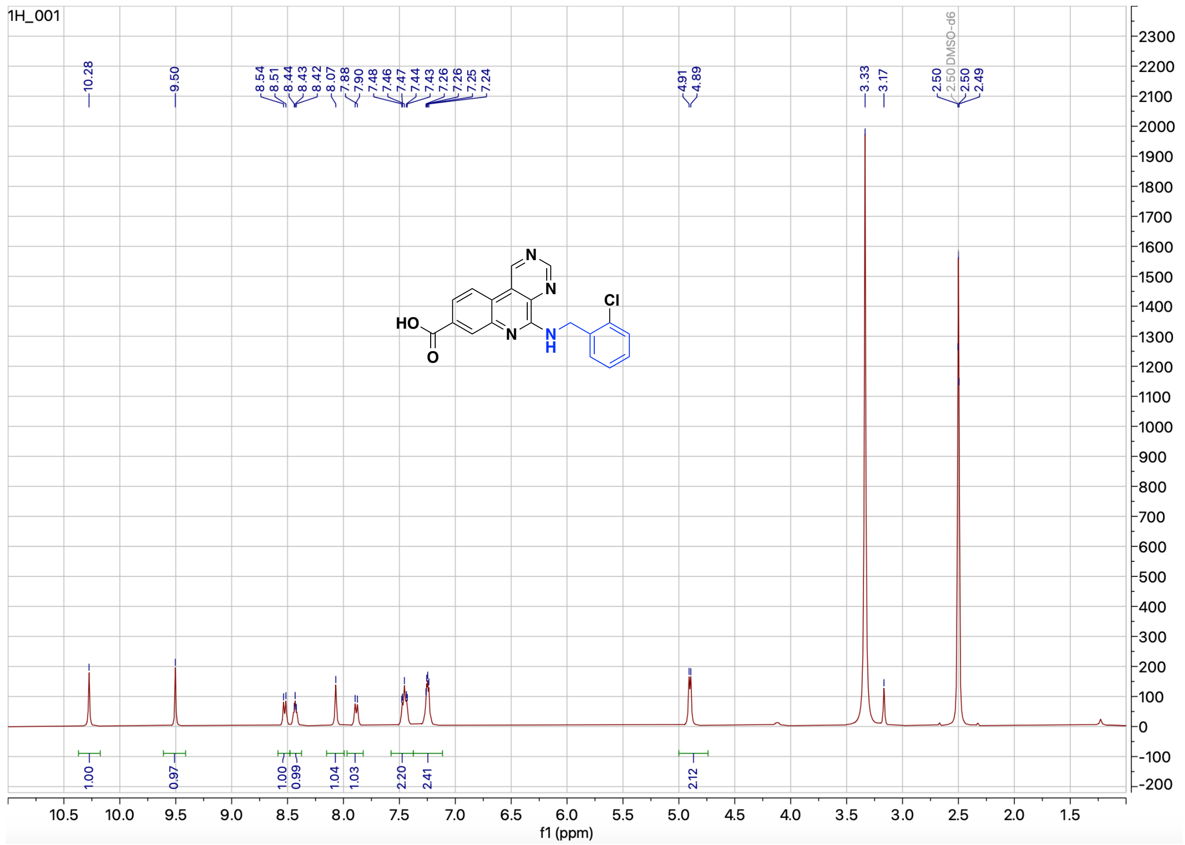

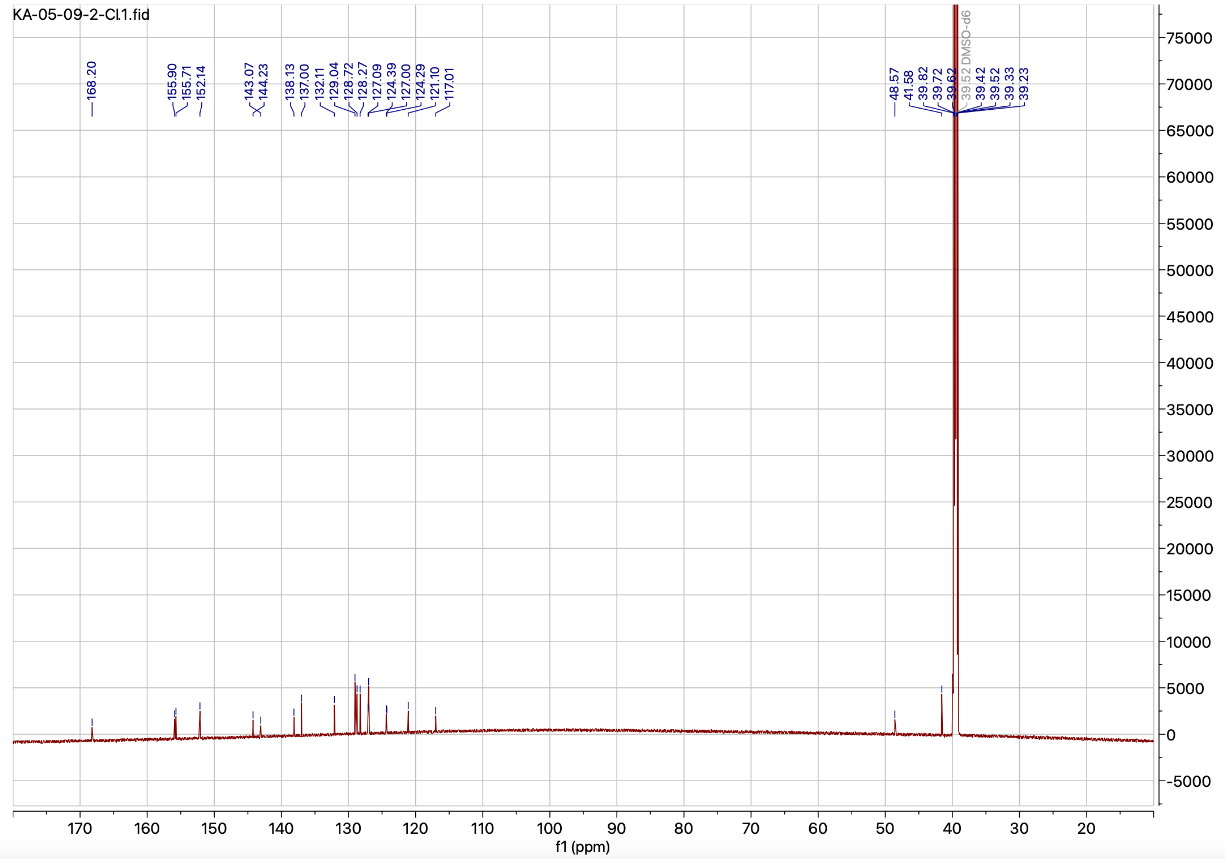

**Figure S3.** $^{1}H$ and $^{13}C$ NMR spectrum of compound **4a**

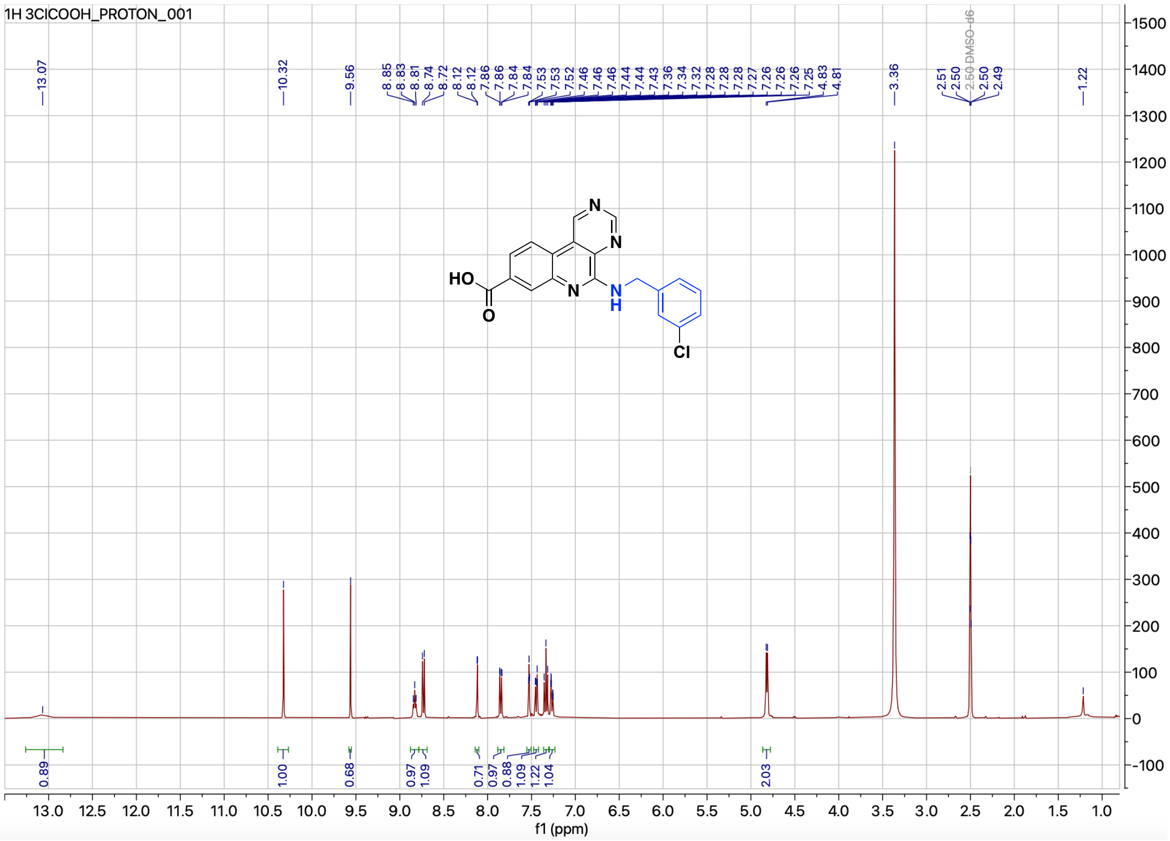

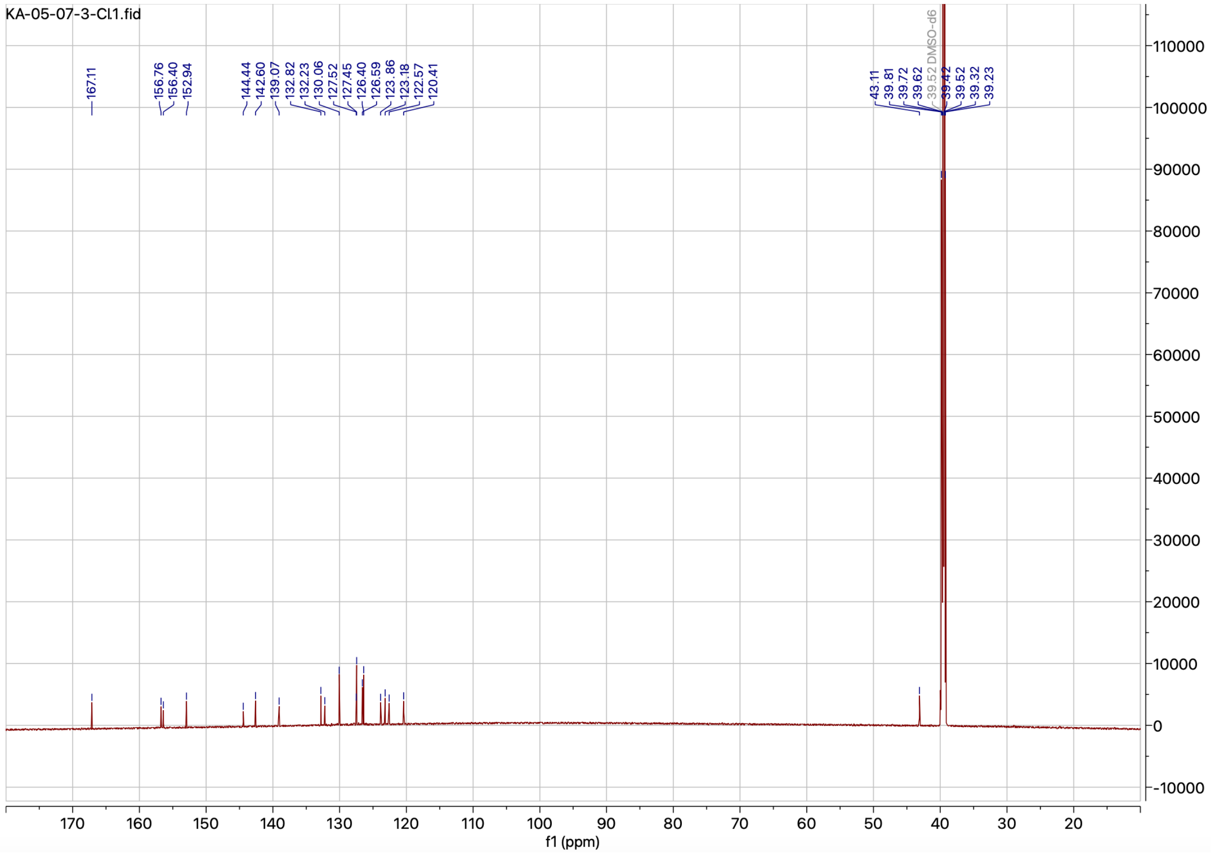


**Figure S4.** $^{1}$H and $^{13}$C NMR spectrum of compound **4b**

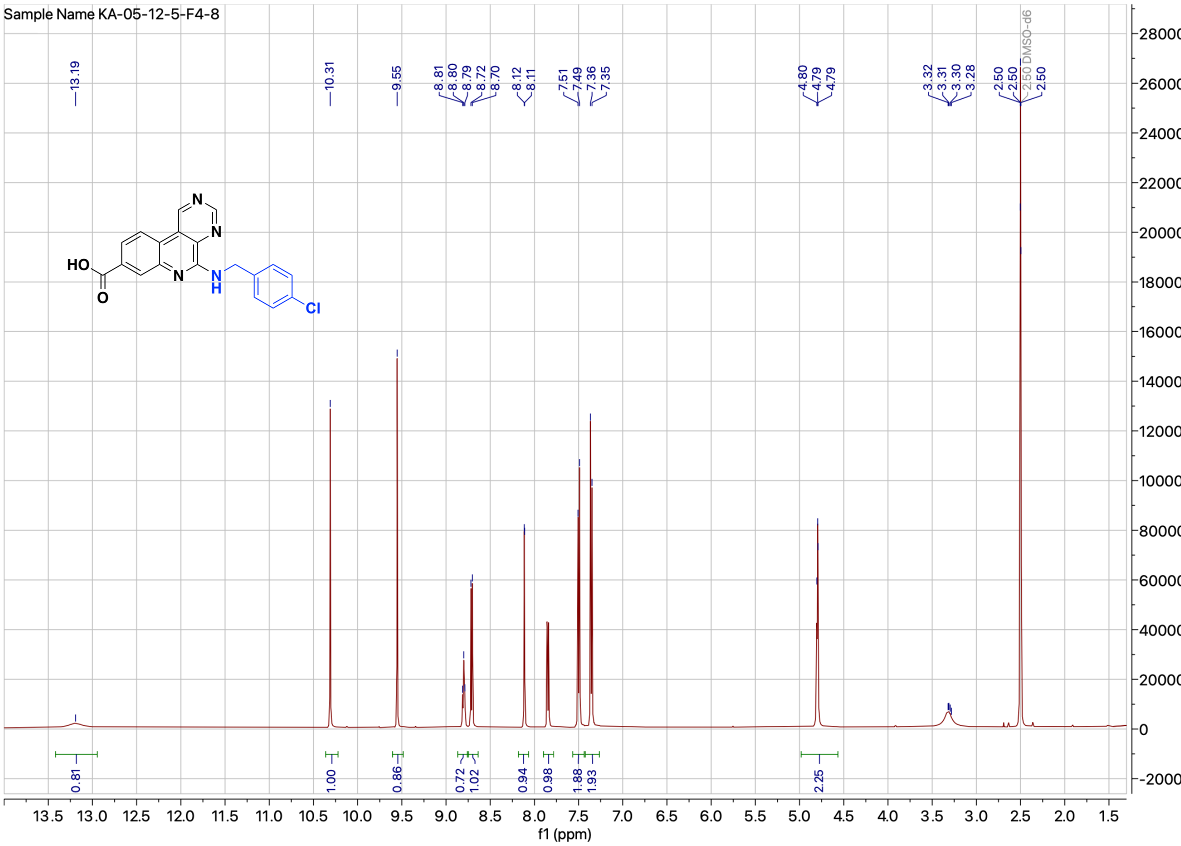


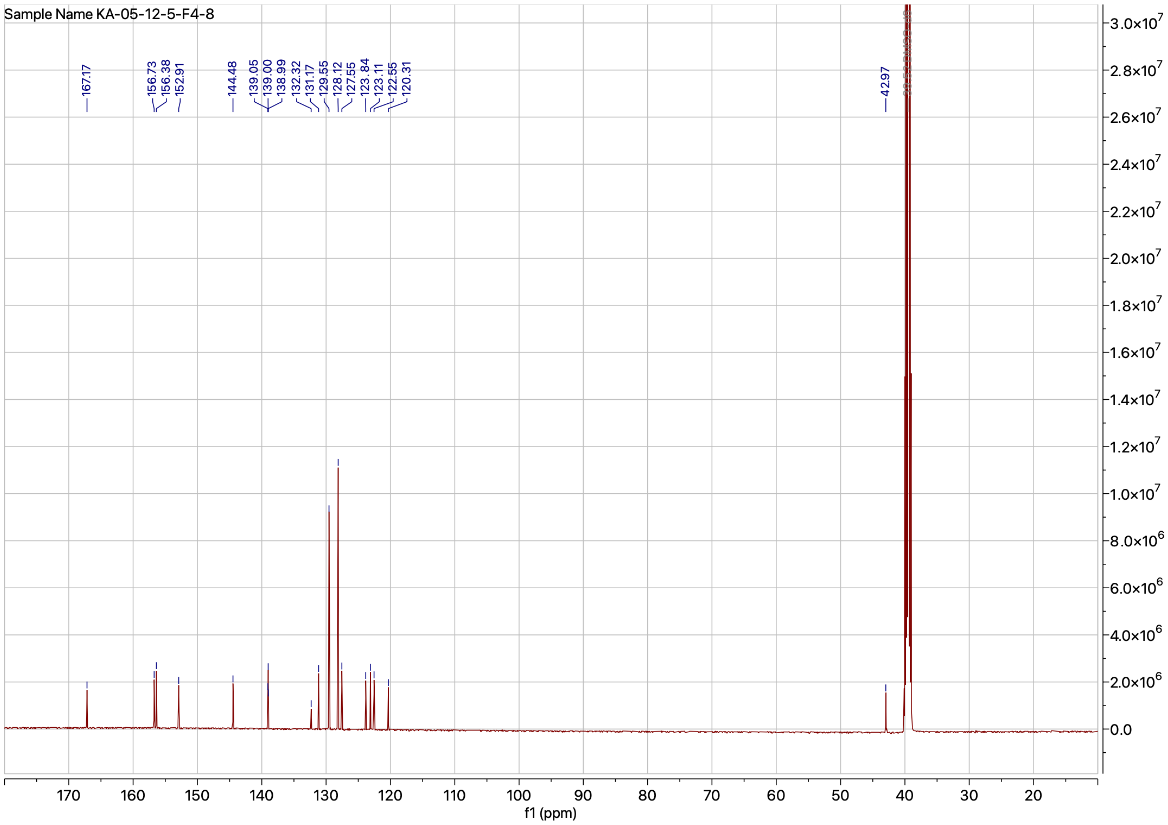


**Figure S5.** $^{1}$H and $^{13}$C NMR spectrum of compound **4c**

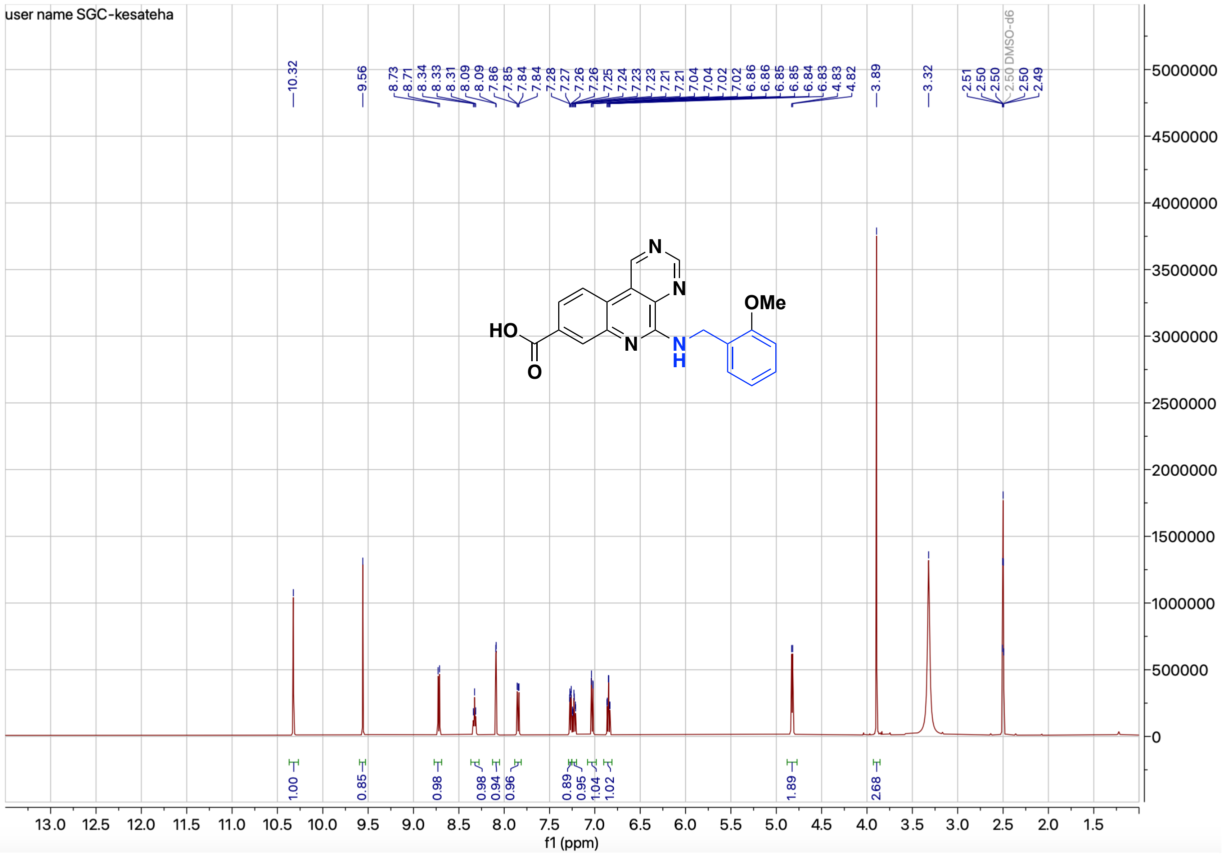


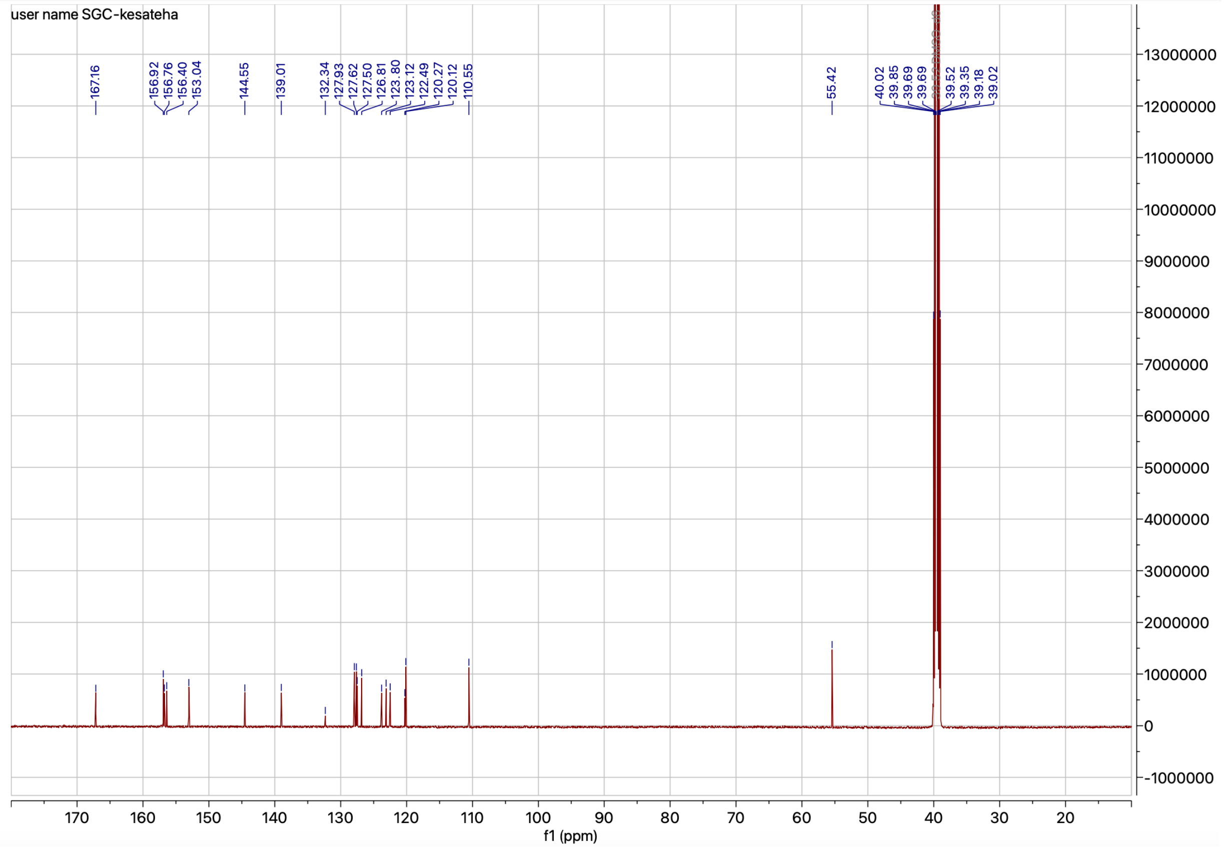


**Figure S6.** [1]H and [13]C NMR spectrum of compound **4d**

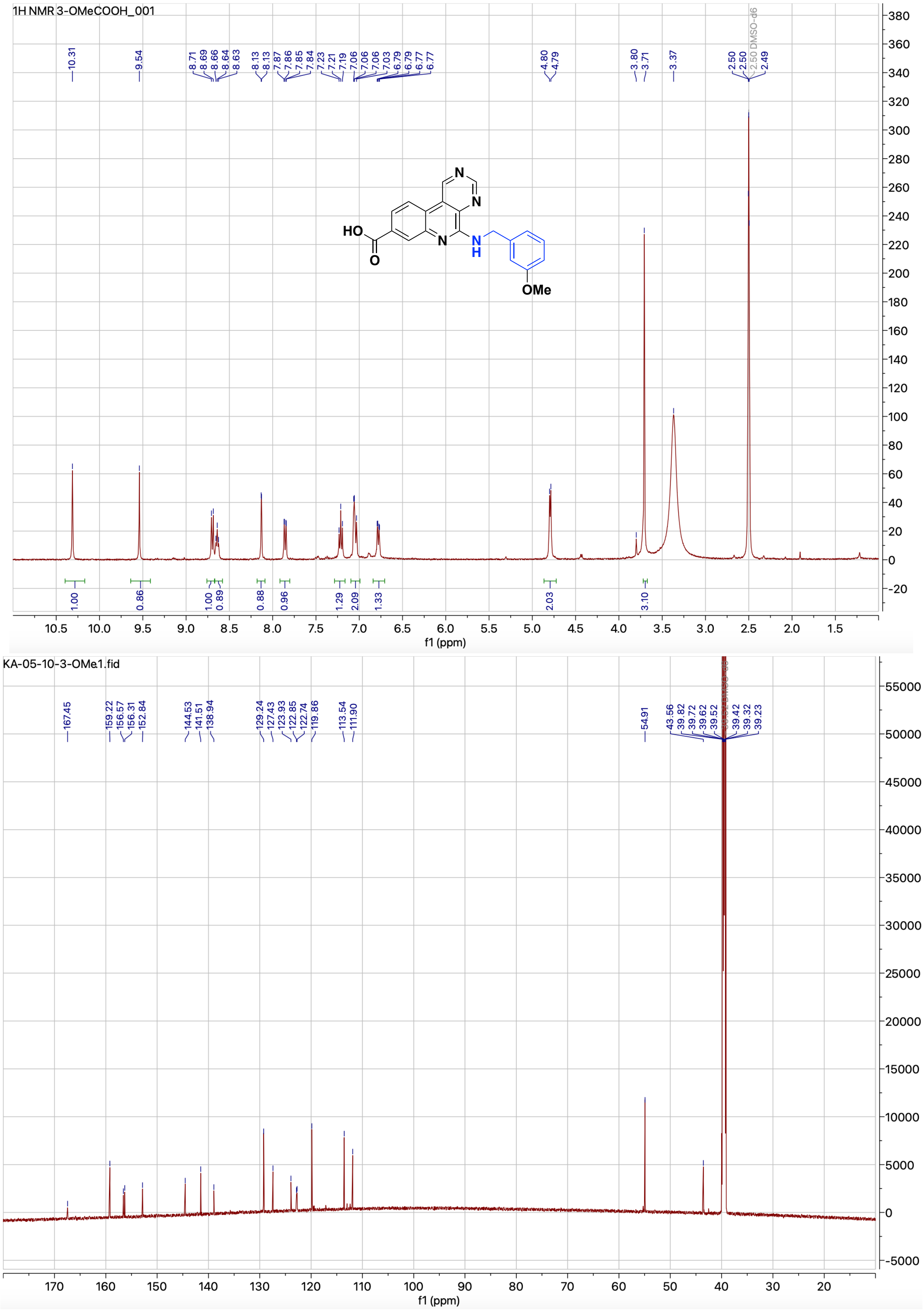


**Figure S7.** $^1$H and $^{13}$C NMR spectrum of compound **4e**

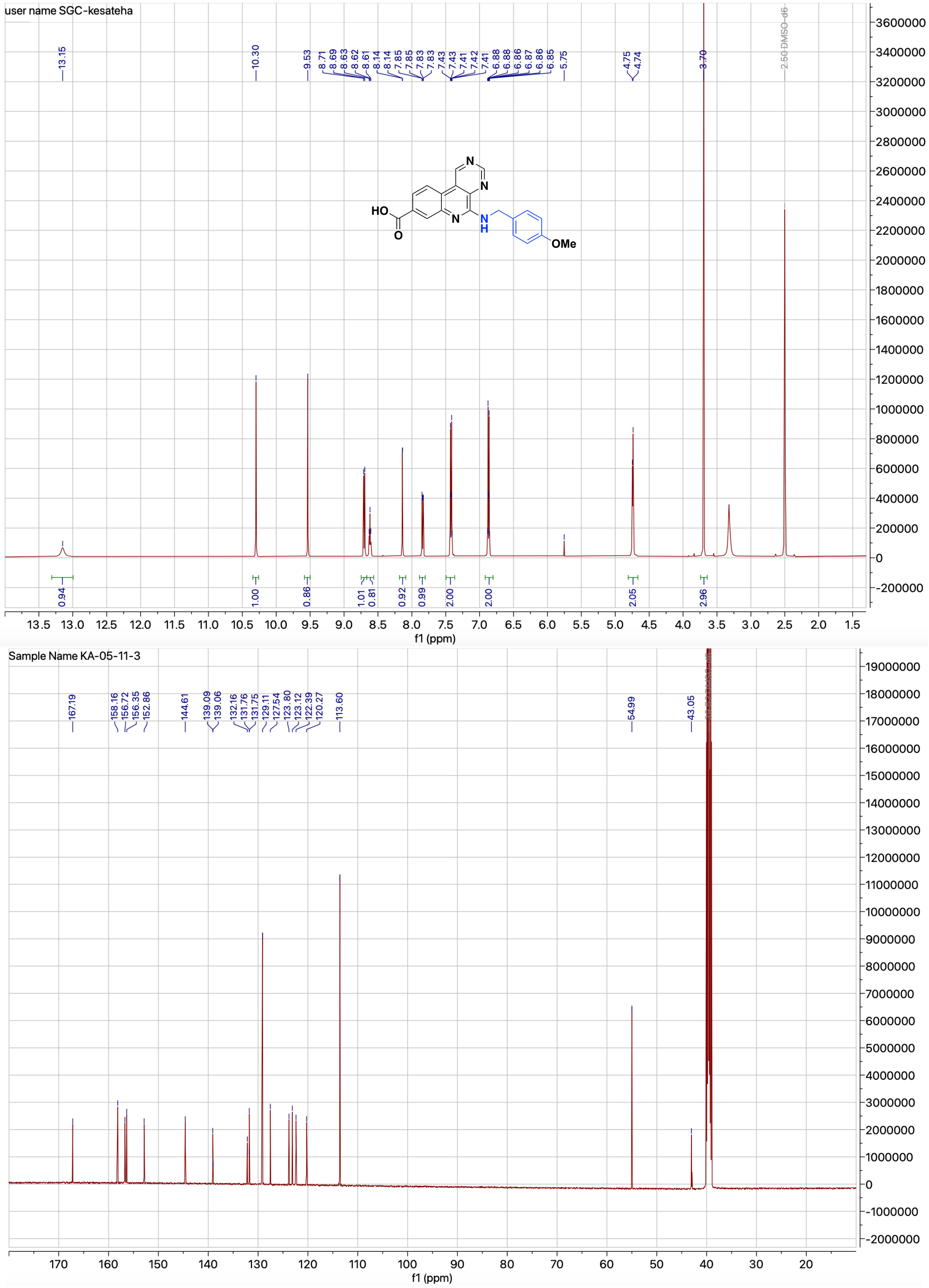


**Figure S8.** $^{1}$H and $^{13}$C NMR spectrum of compound **4f**

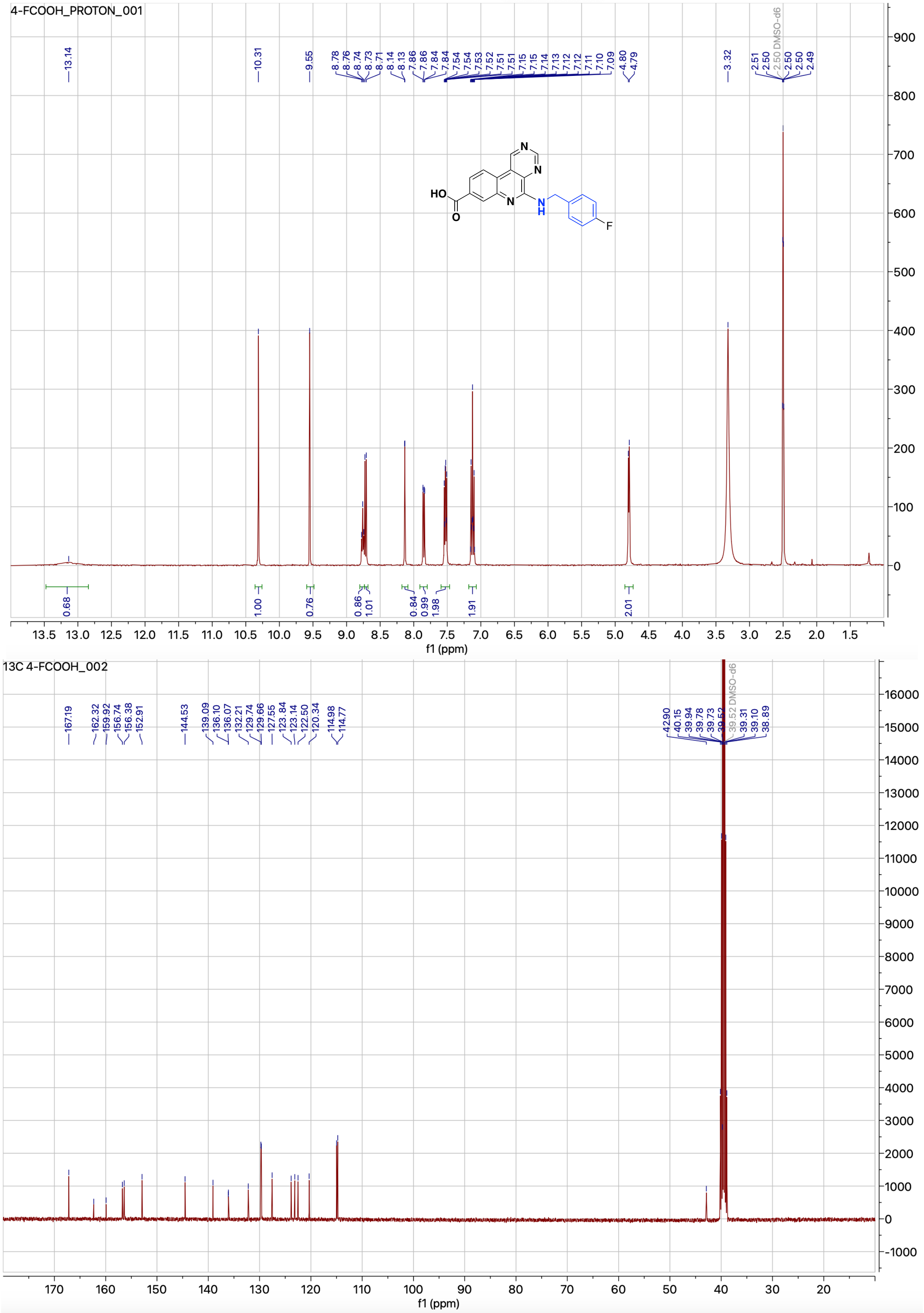

**Figure S9.** [1]H and [13]C NMR spectrum of compound **4g**

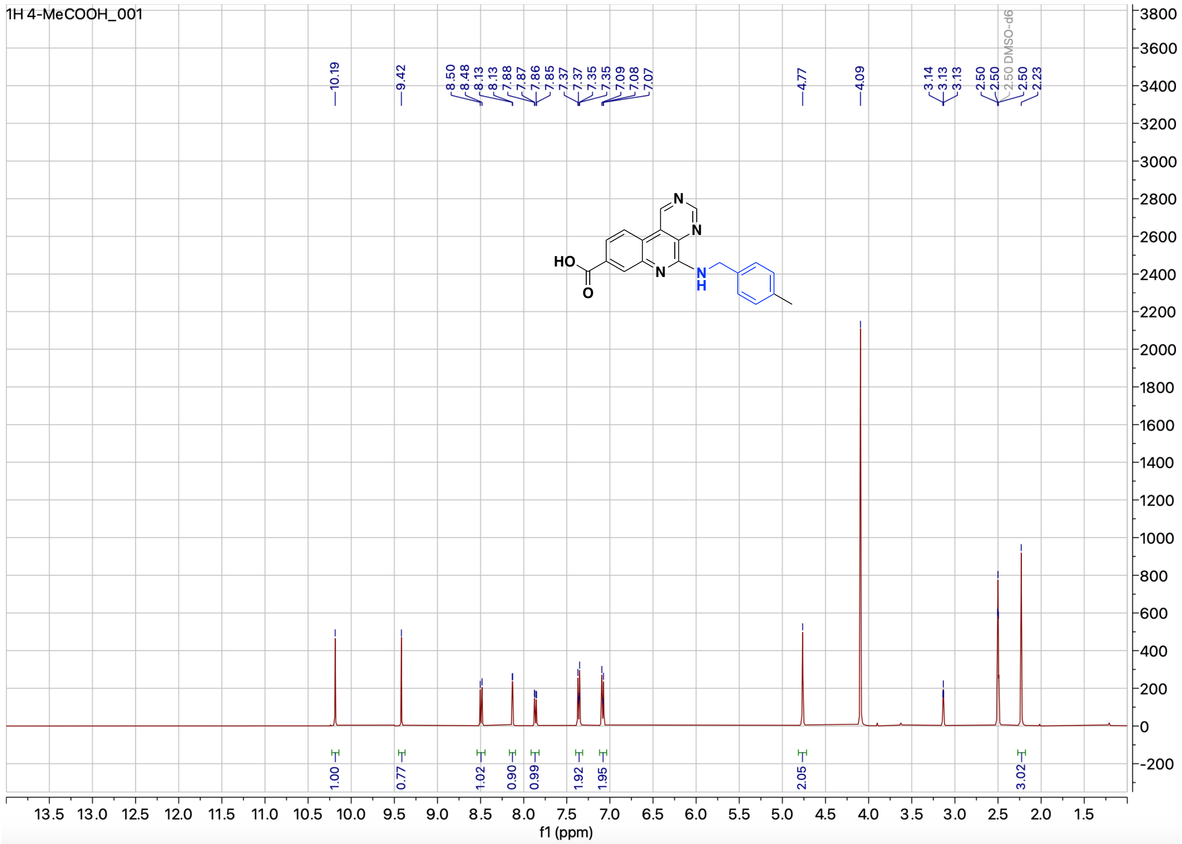


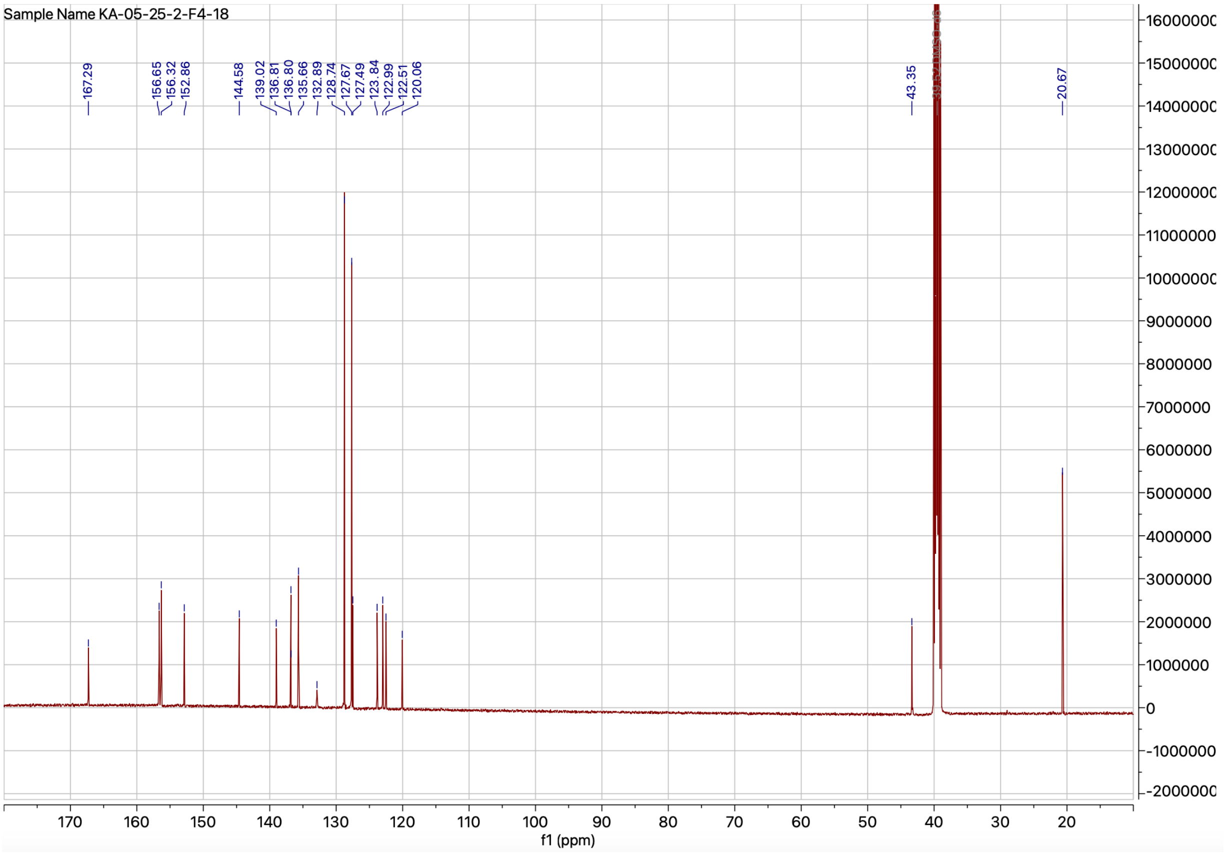


**Figure S10.** $^{1}$H and $^{13}$C NMR spectrum of compound **4h**

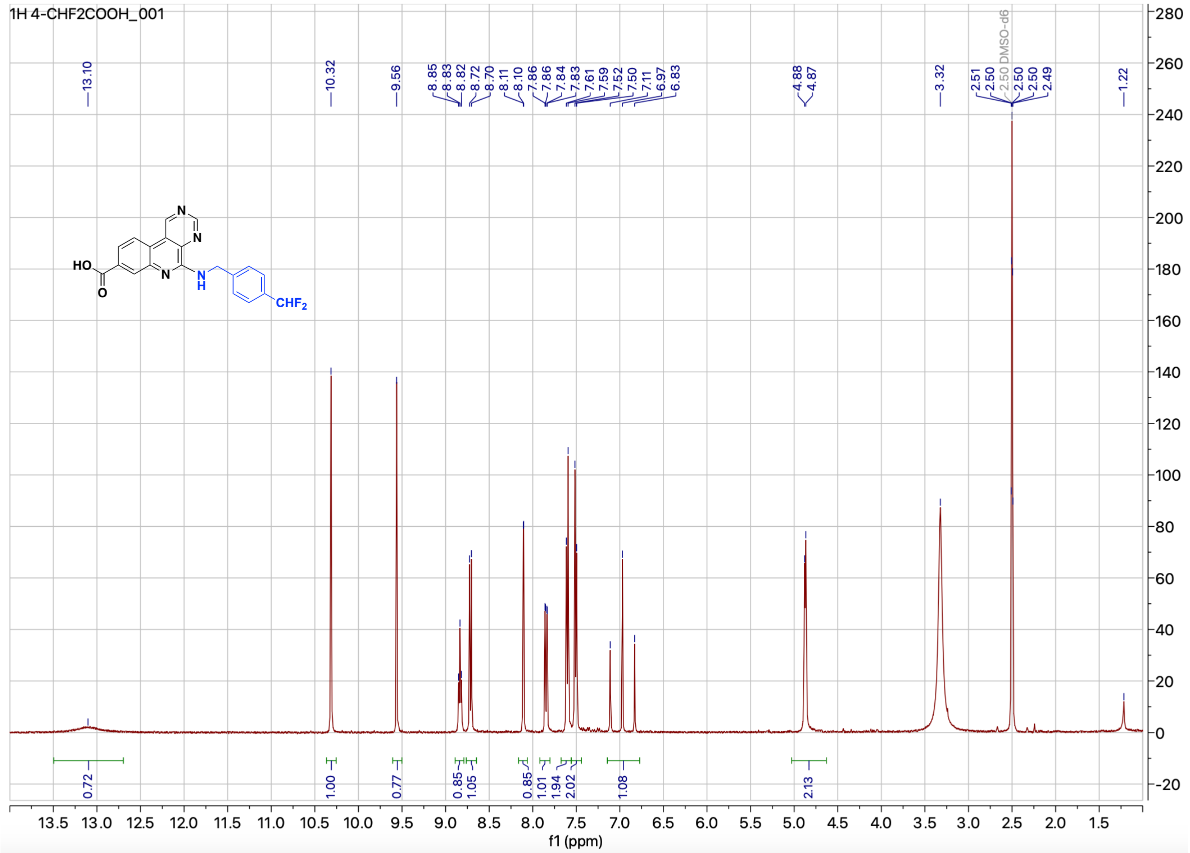

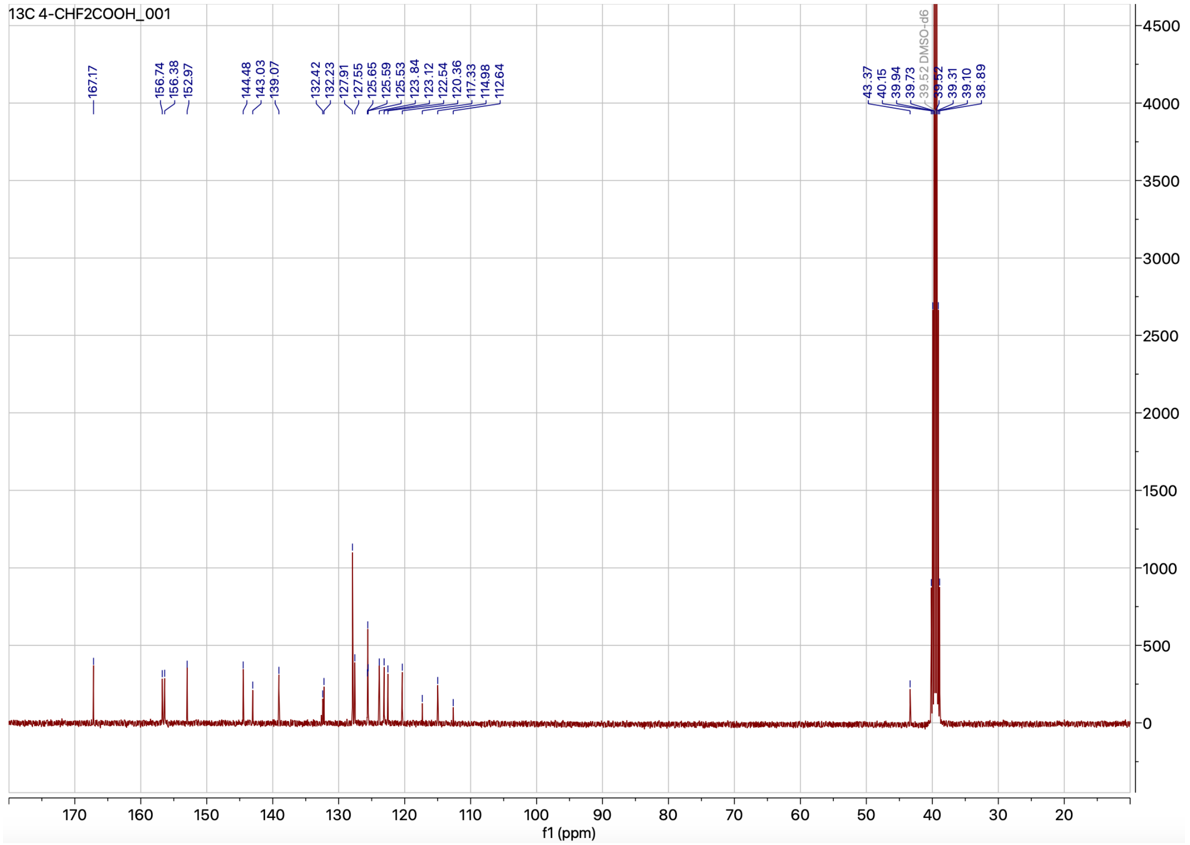

**Figure S11.** ¹H and ¹³C NMR spectrum of compound **4i**

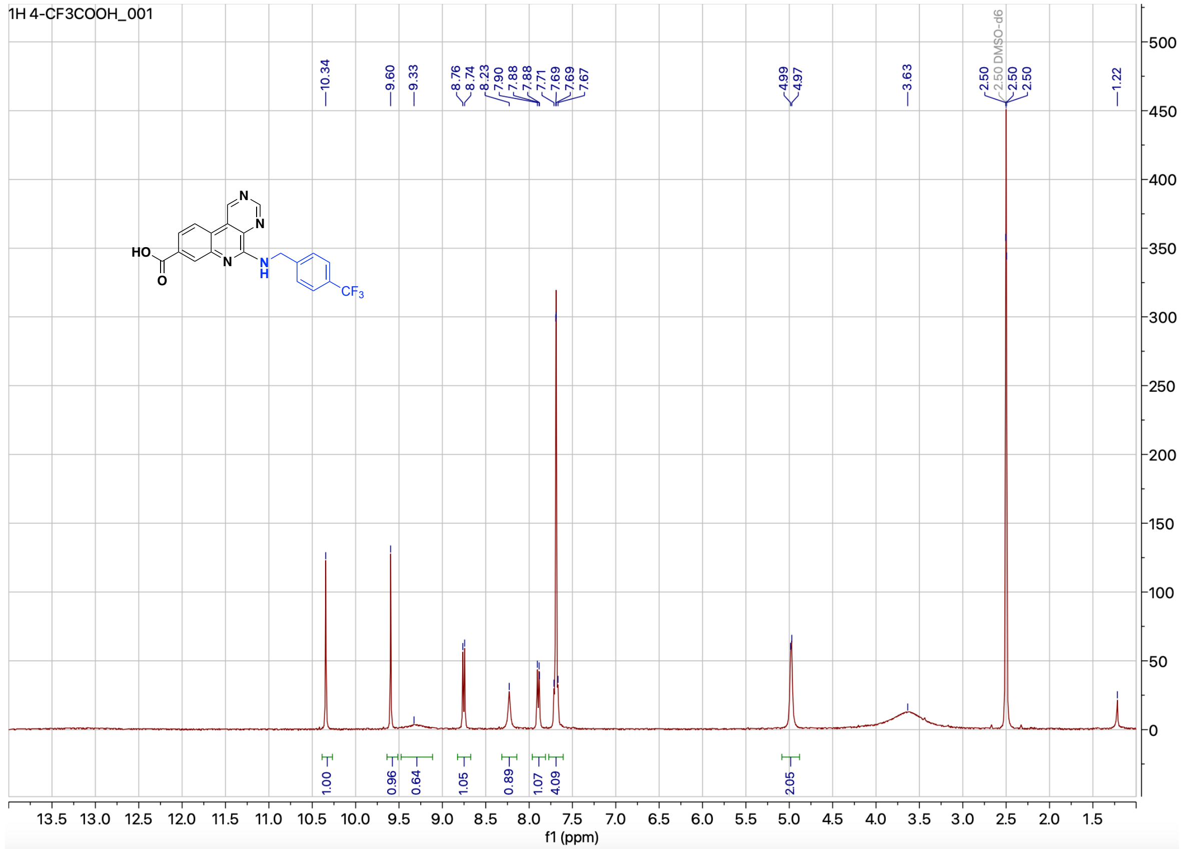

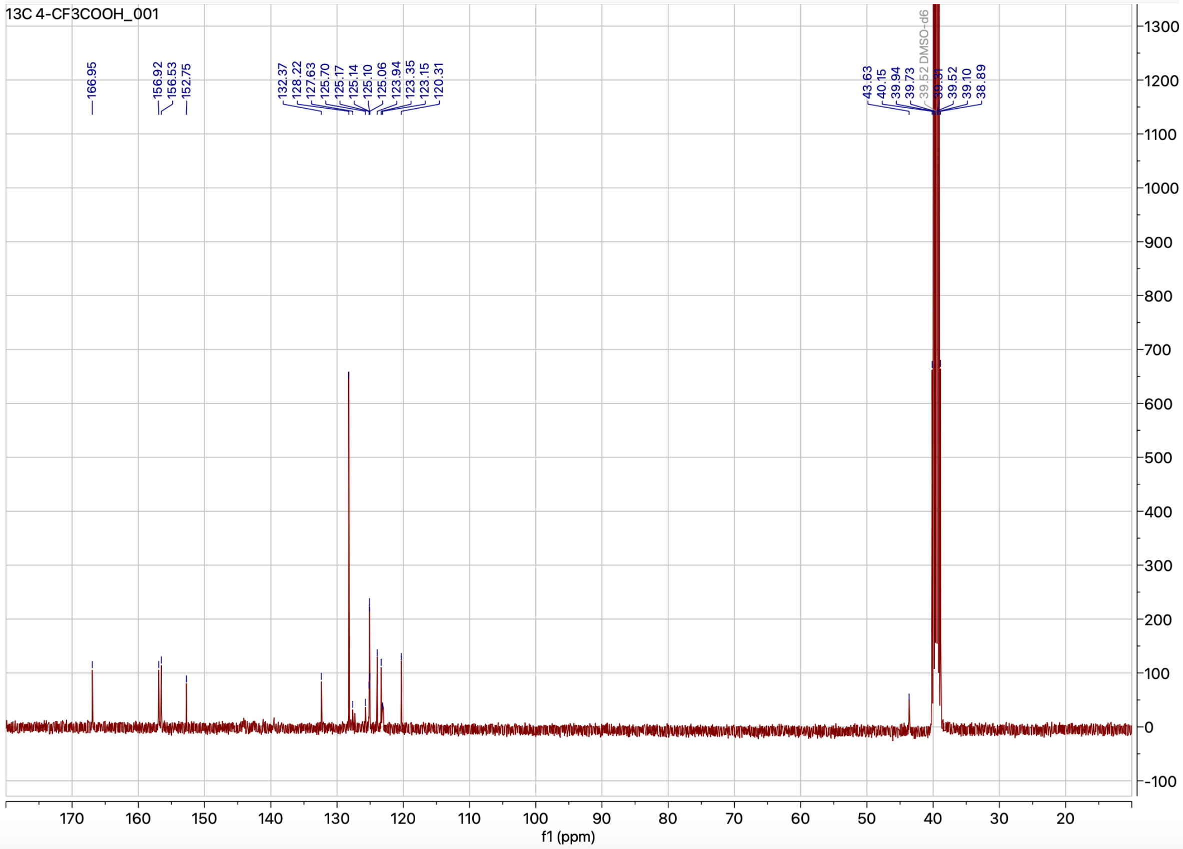

**Figure S12.** [1]H and [13]C NMR spectrum of compound **4j**

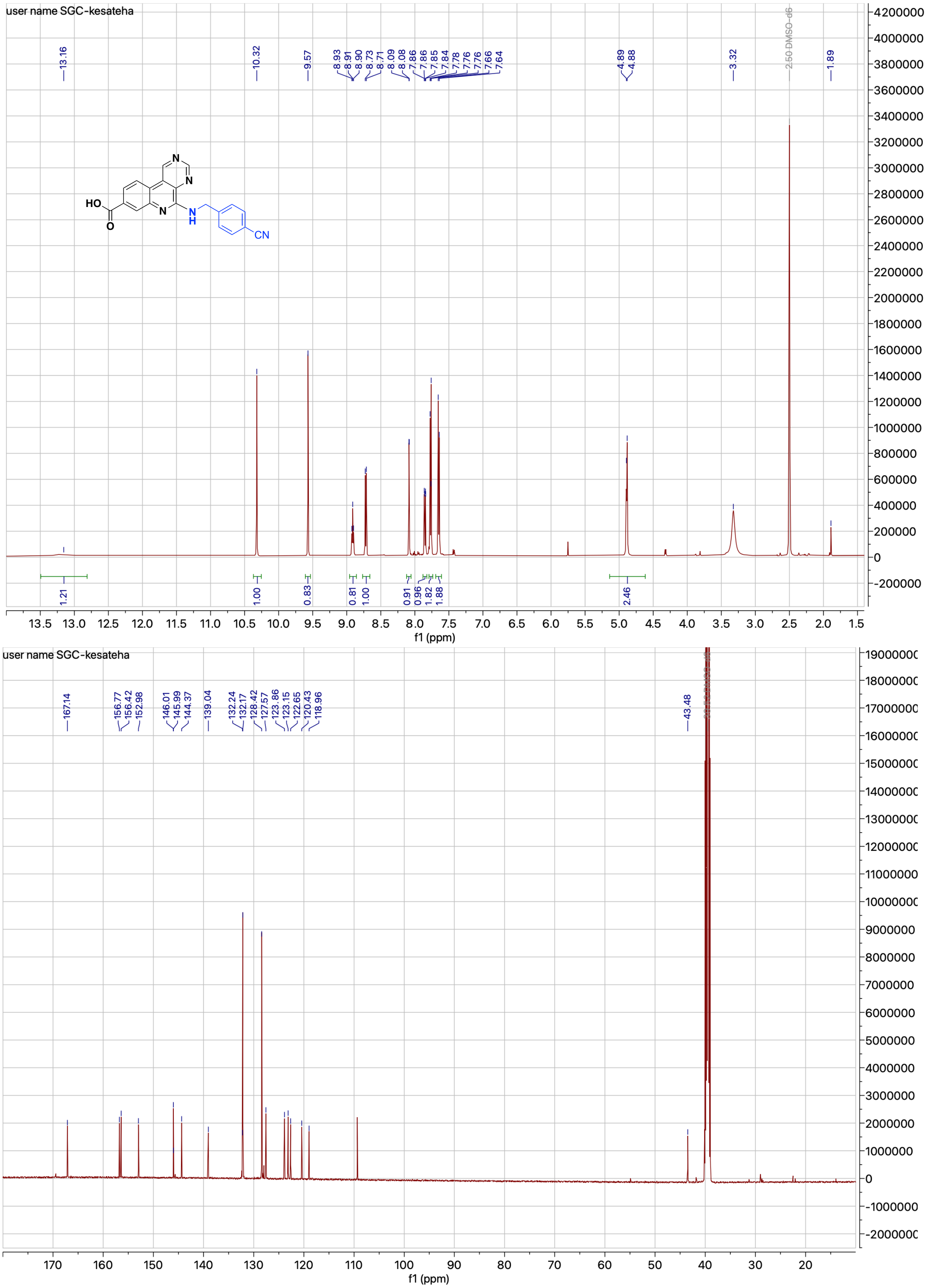


**Figure S13.** $^{1}$H and $^{13}$C NMR spectrum of compound **4k**

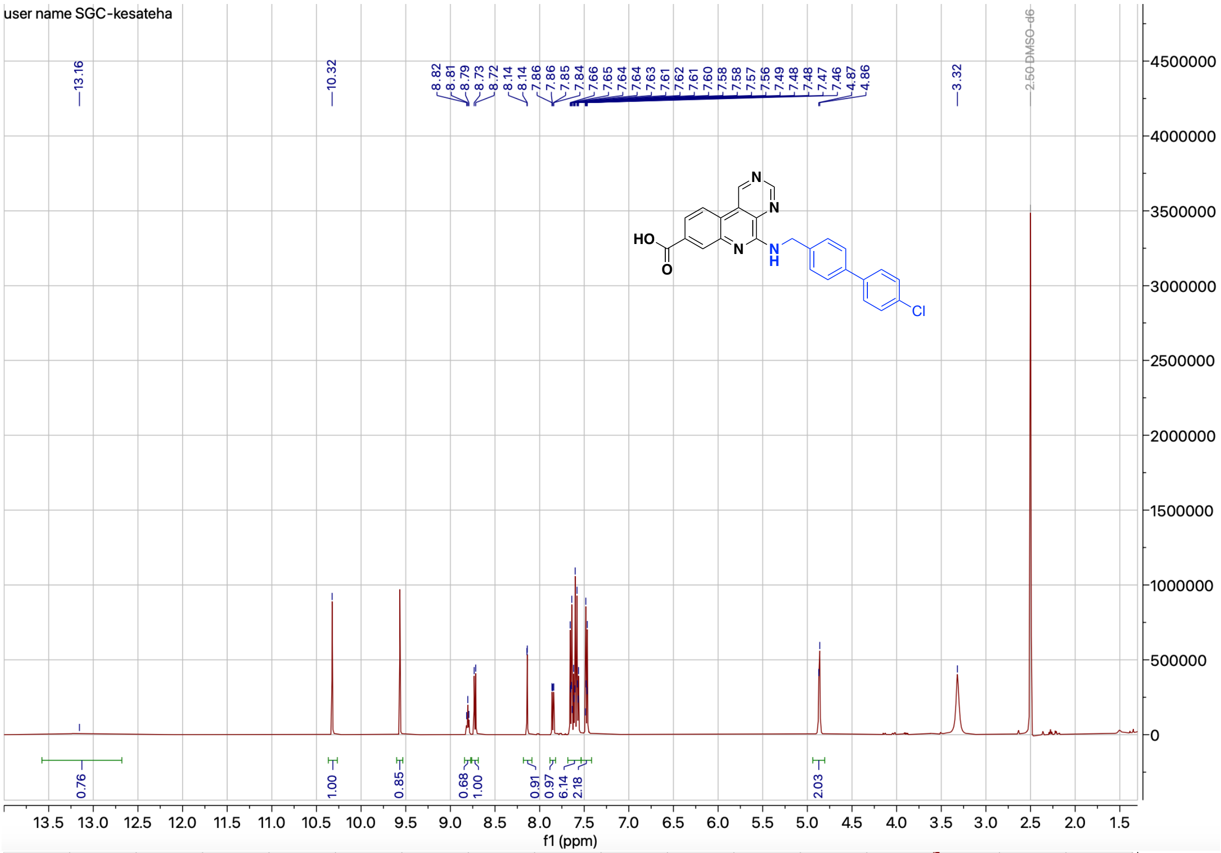


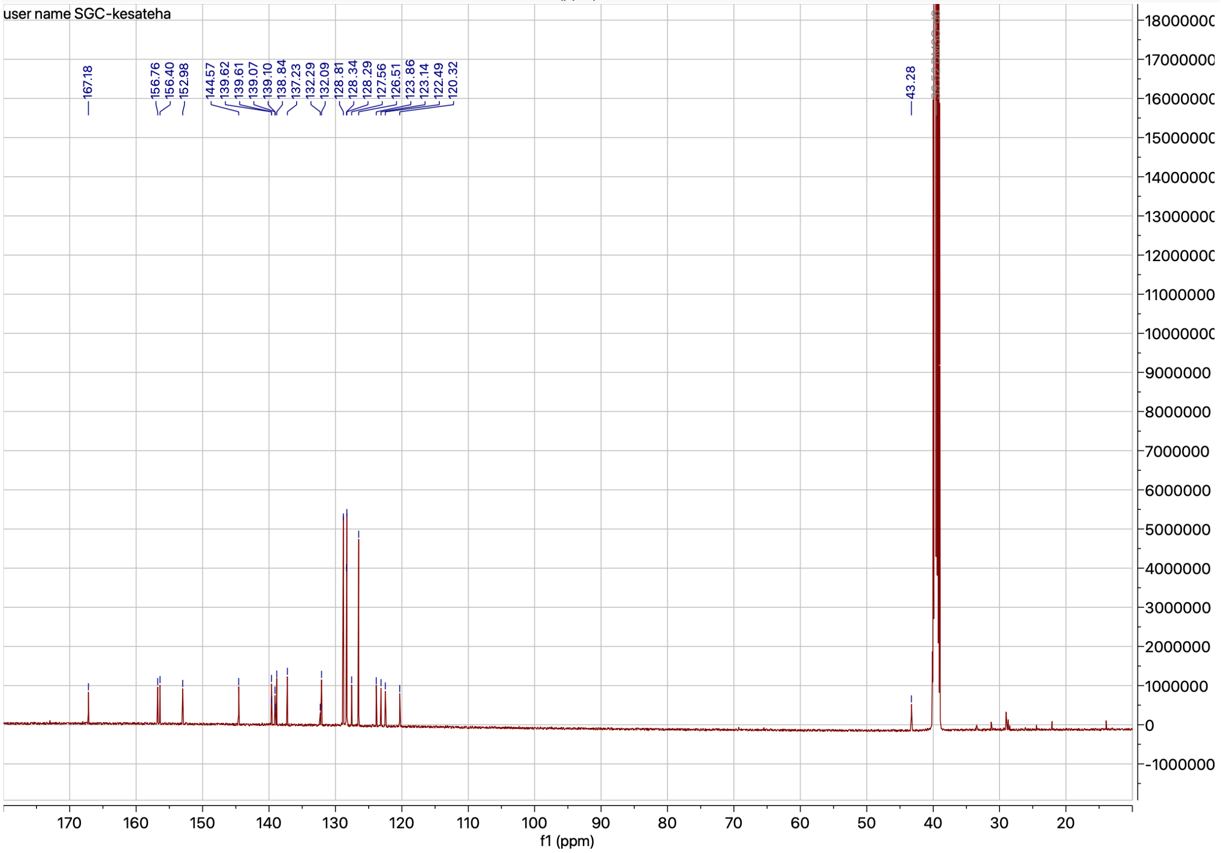


**Figure S14.** $^{1}$H and $^{13}$C NMR spectrum of compound **4l**

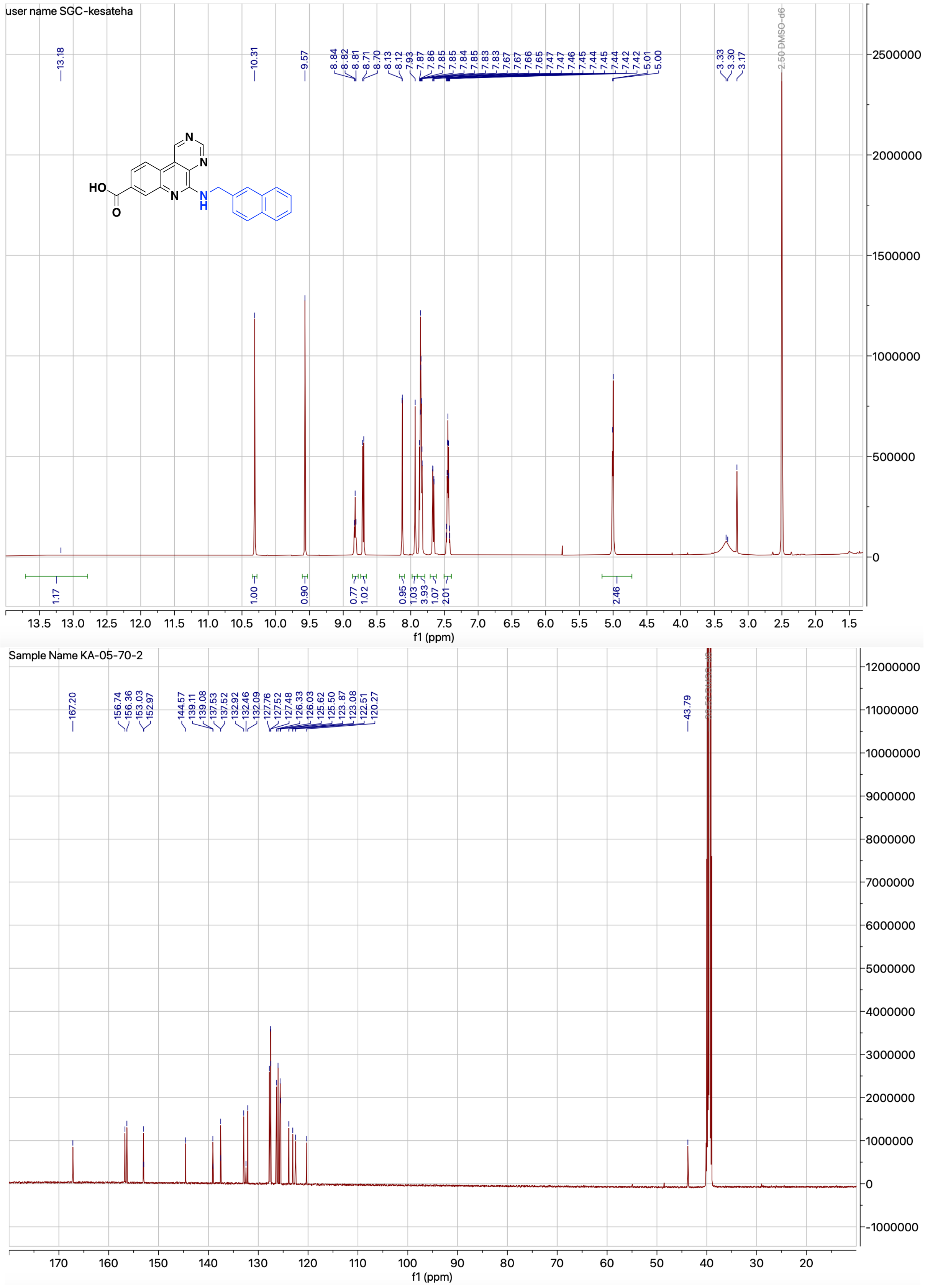

**Figure S15.** $^{1}$H and $^{13}$C NMR spectrum of compound **4m**

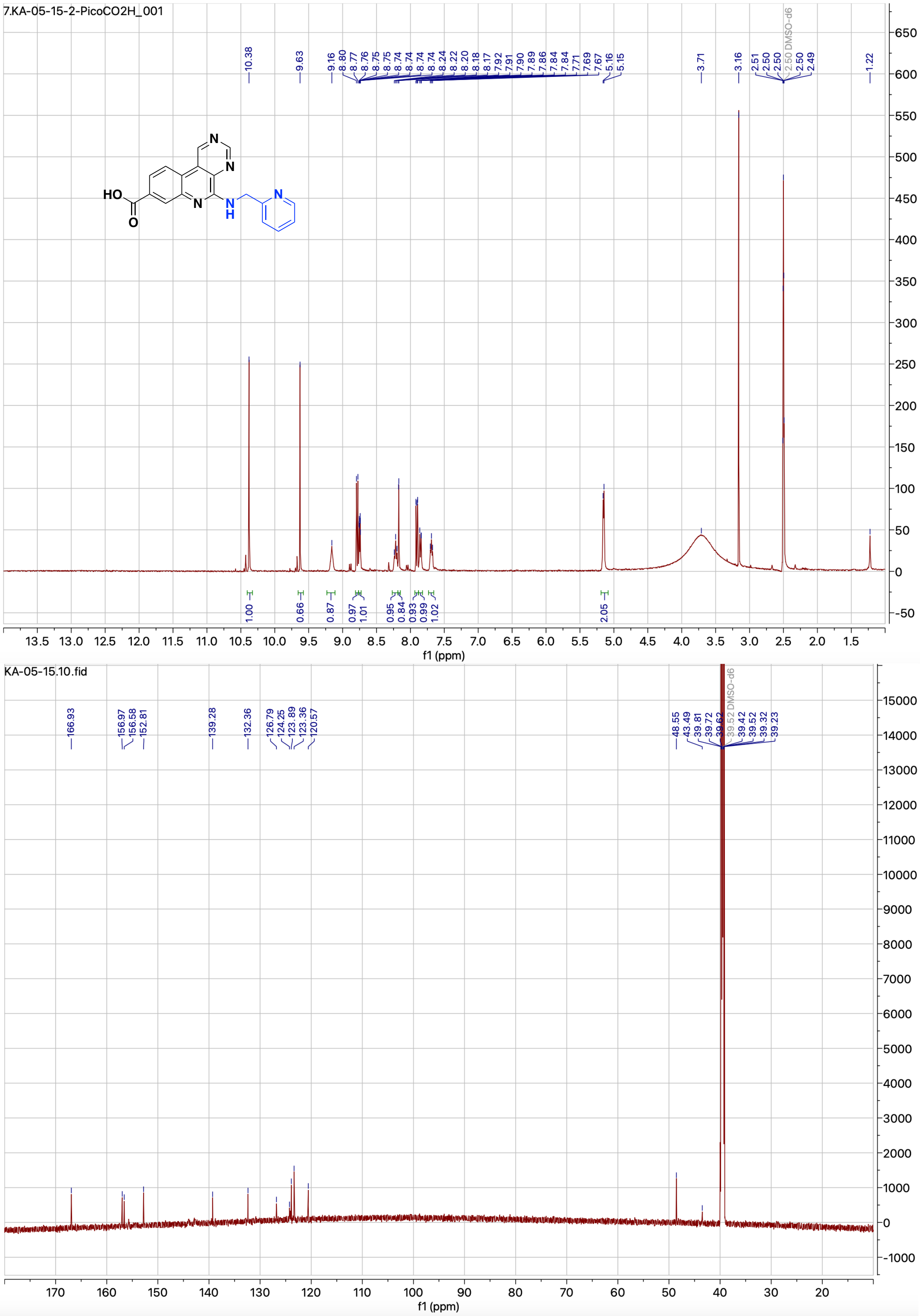


**Figure S16.** $^{1}$H and $^{13}$C NMR spectrum of compound **4n**

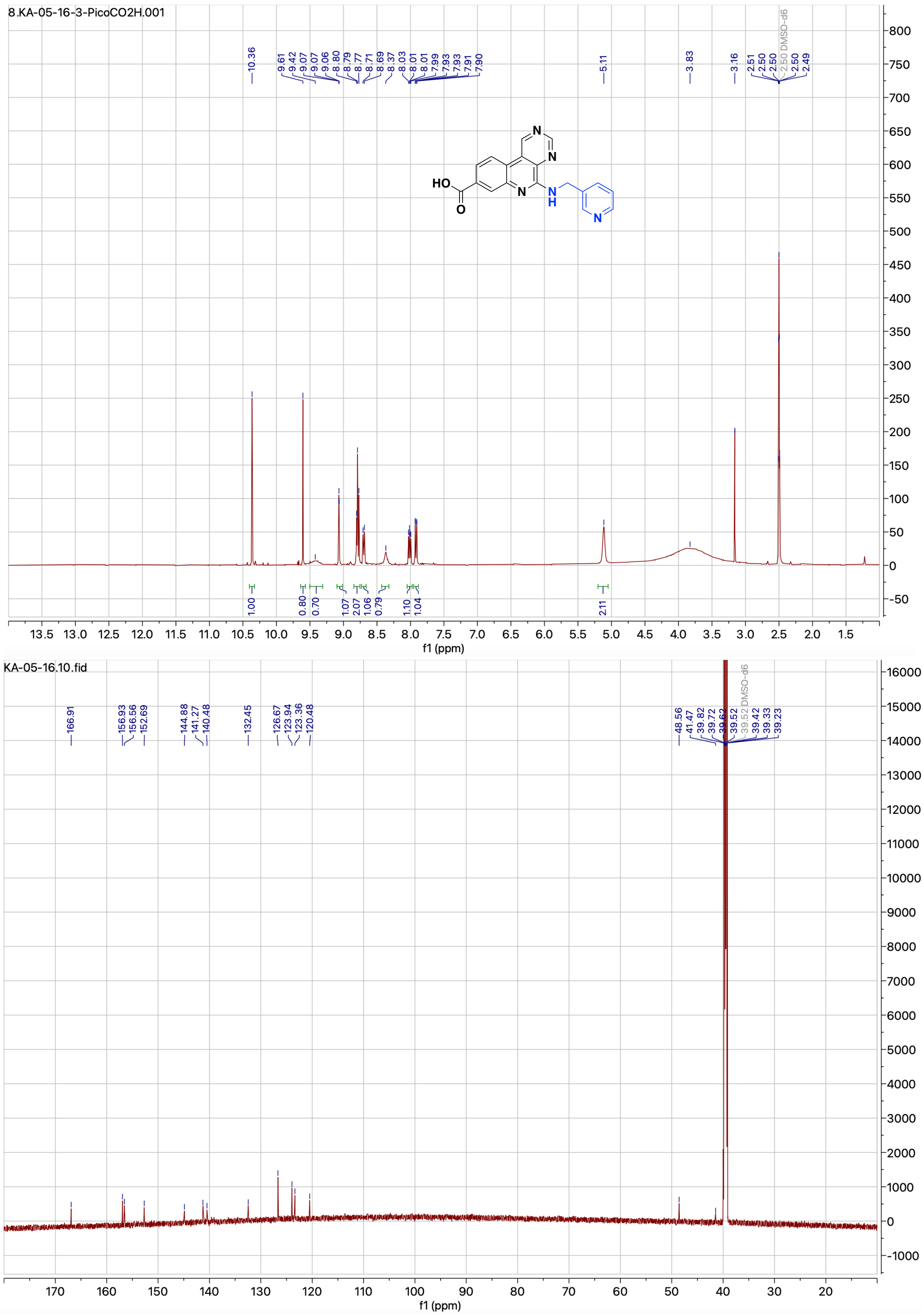


**Figure S17.** $^{1}$H and $^{13}$C NMR spectrum of compound **4o**

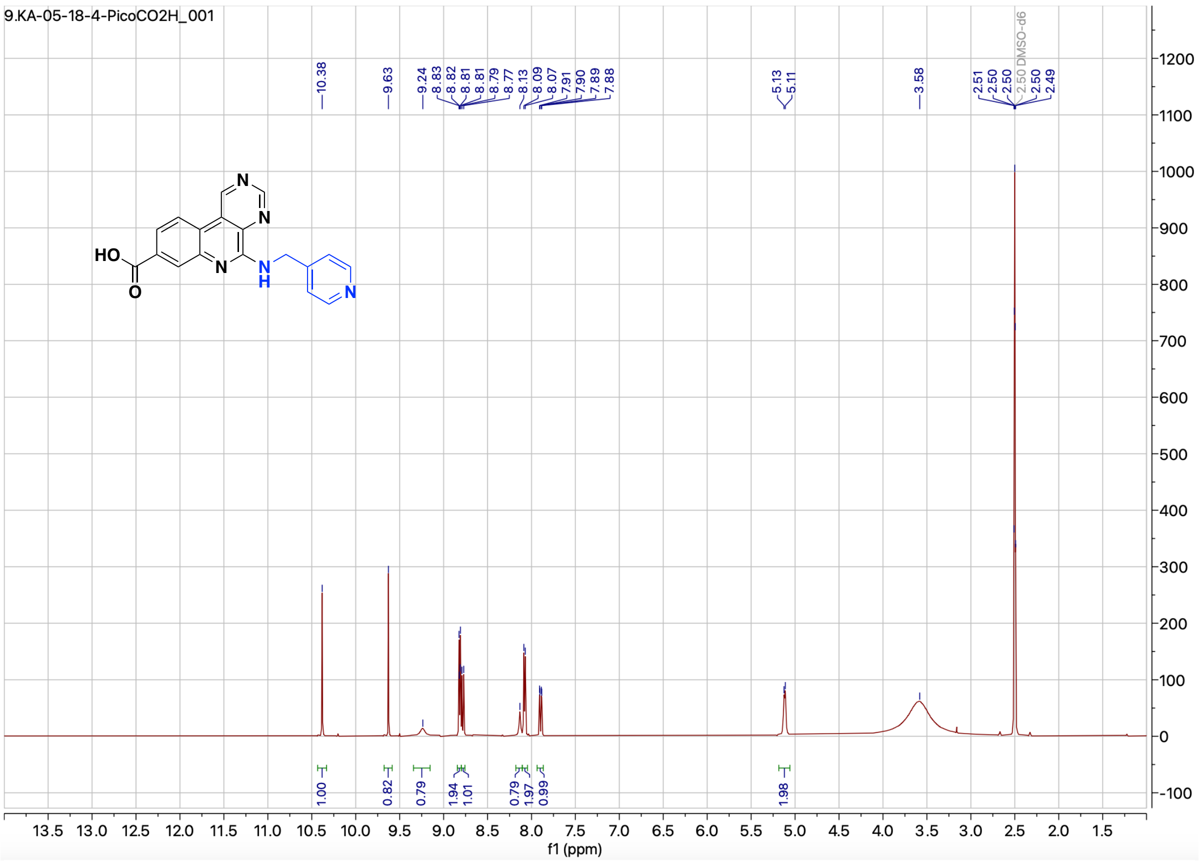


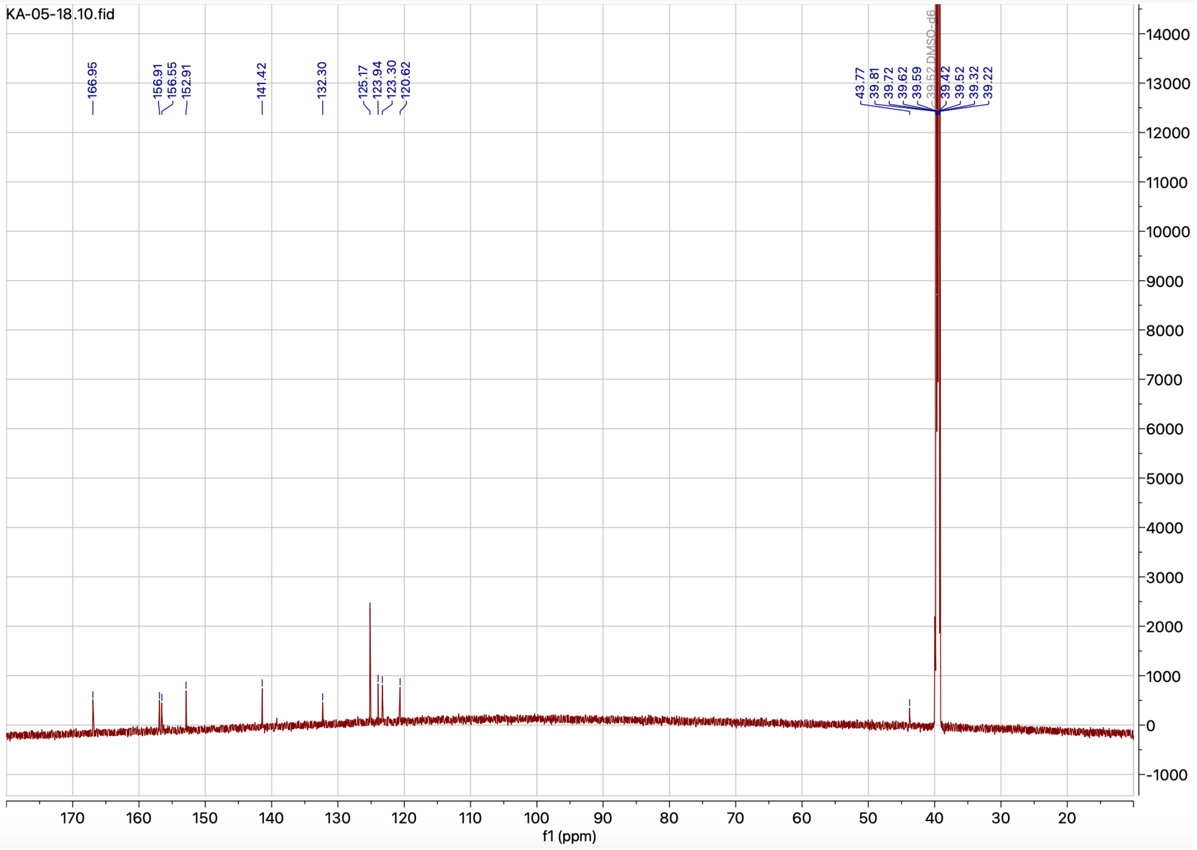


**Figure S18.** $^{1}$H and $^{13}$C NMR spectrum of compound **4p**

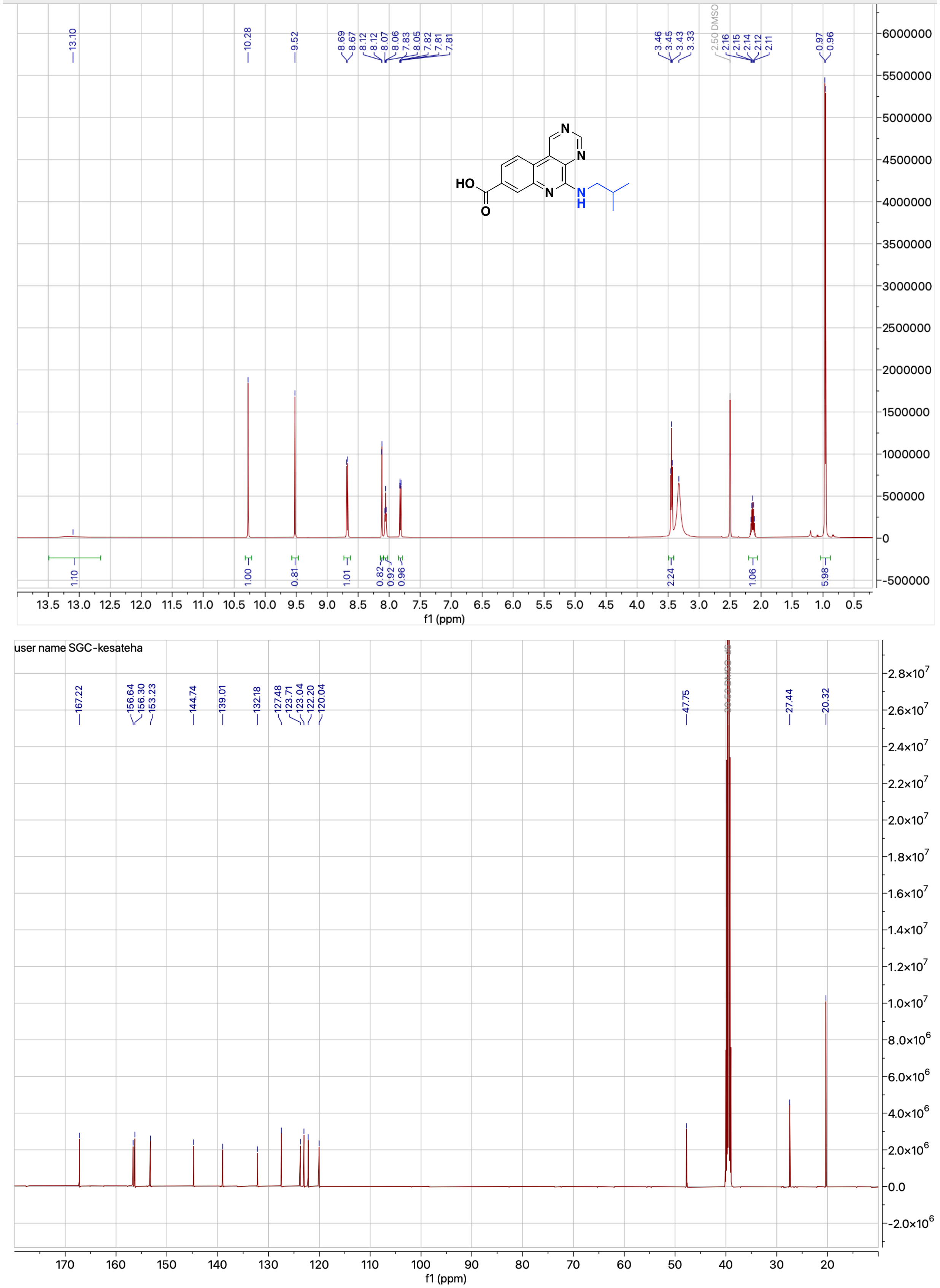


**Figure S19.** $^{1}H$ and $^{13}C$ NMR spectrum of compound **4q**

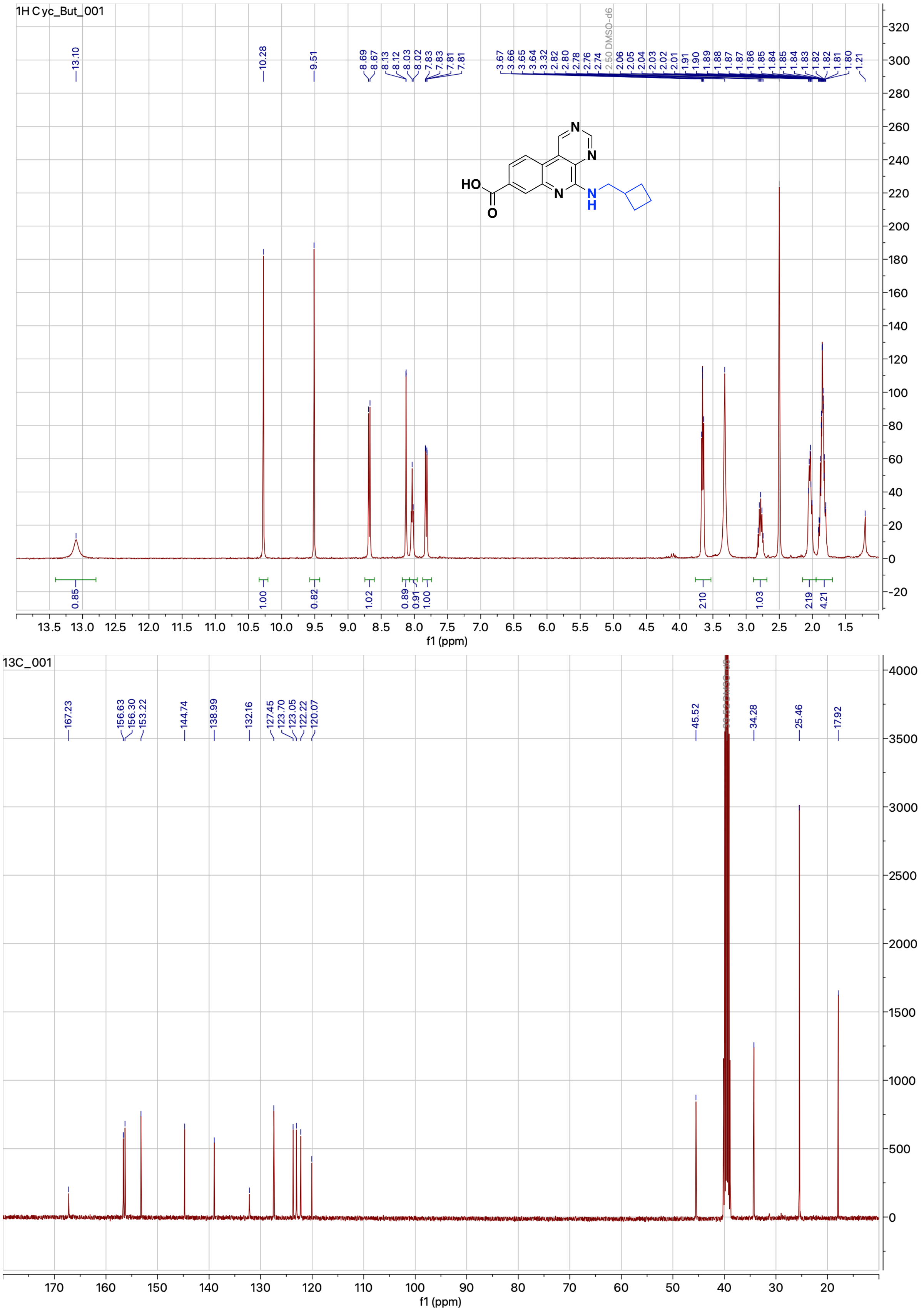


**Figure S20.** $^{1}$H and $^{13}$C NMR spectrum of compound **4r**

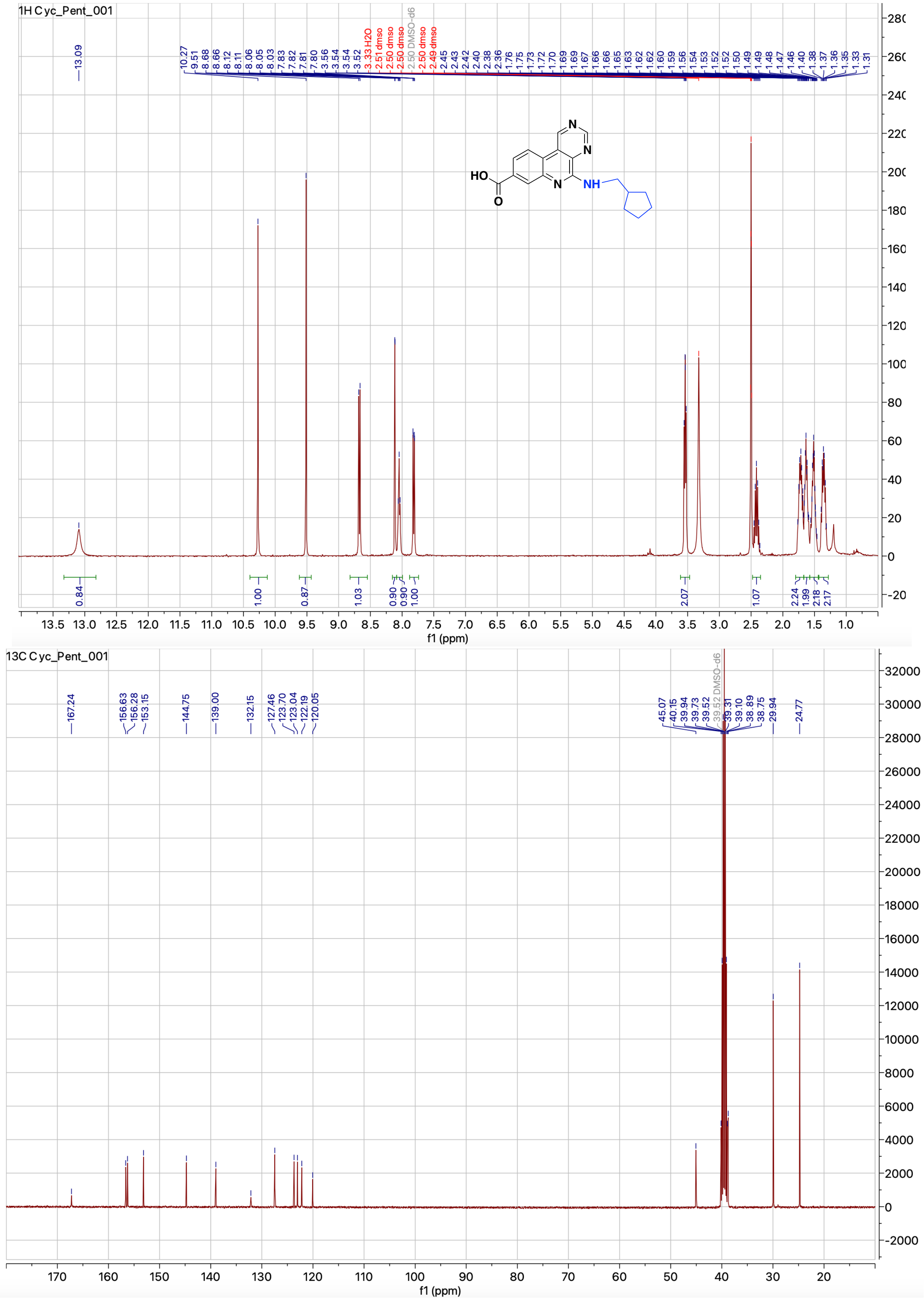

**Figure S21.** $^{1}$H and $^{13}$C NMR spectrum of compound **4s**

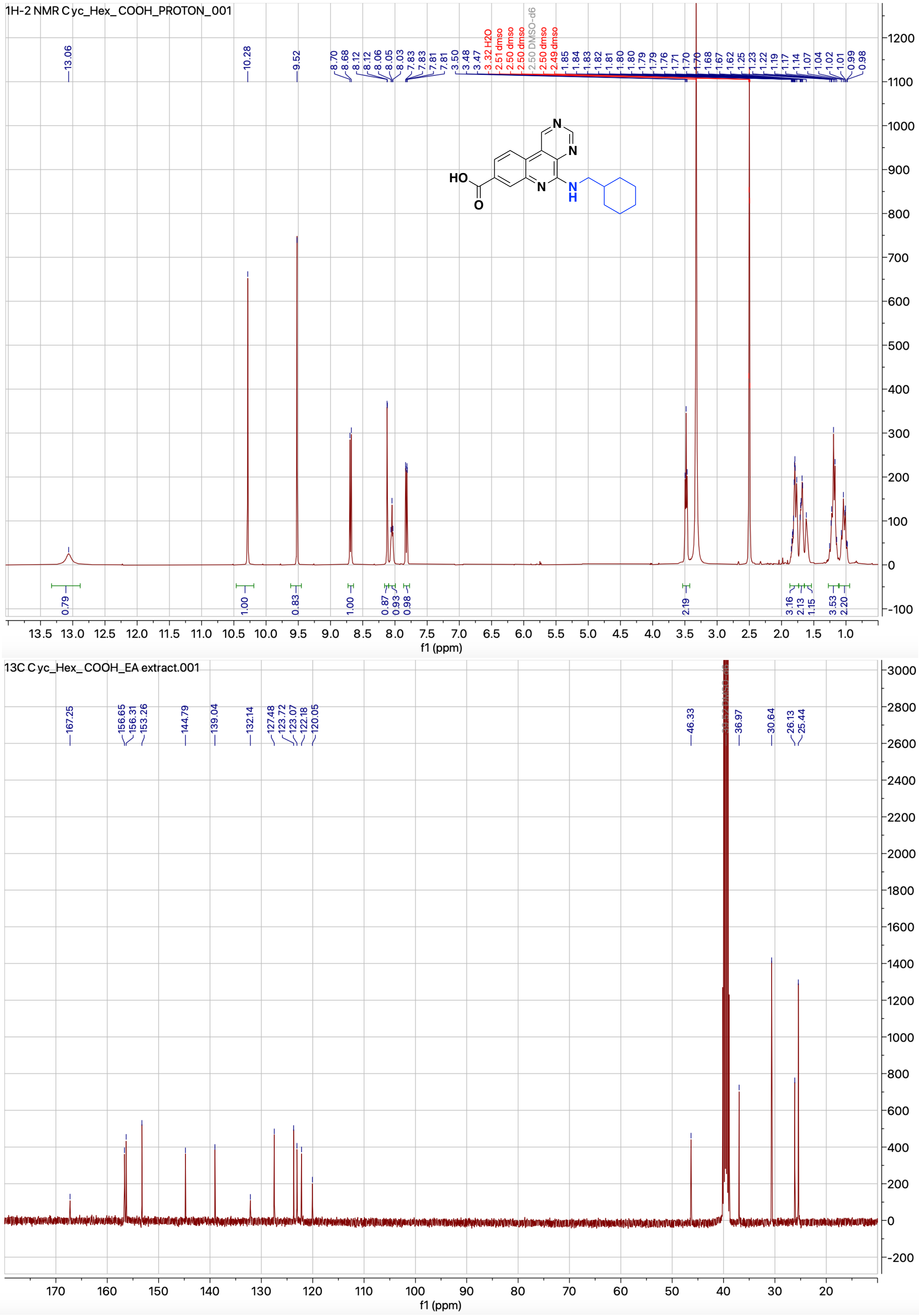


b $^{1}H$ and $^{13}C$ NMR spectrum of compound **4t**

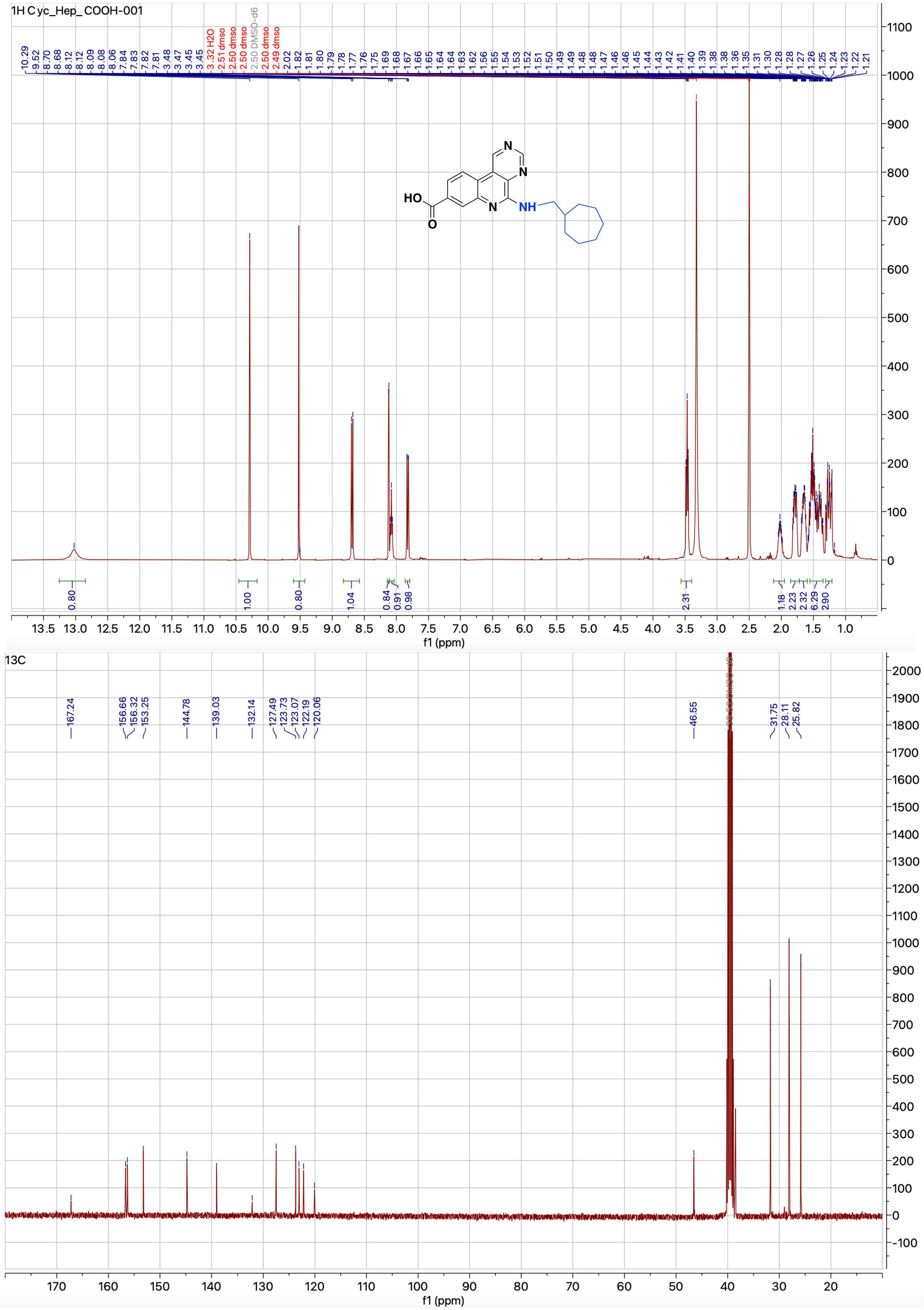


**Figure S23.** $^{1}$H and $^{13}$C NMR spectrum of compound **4u**

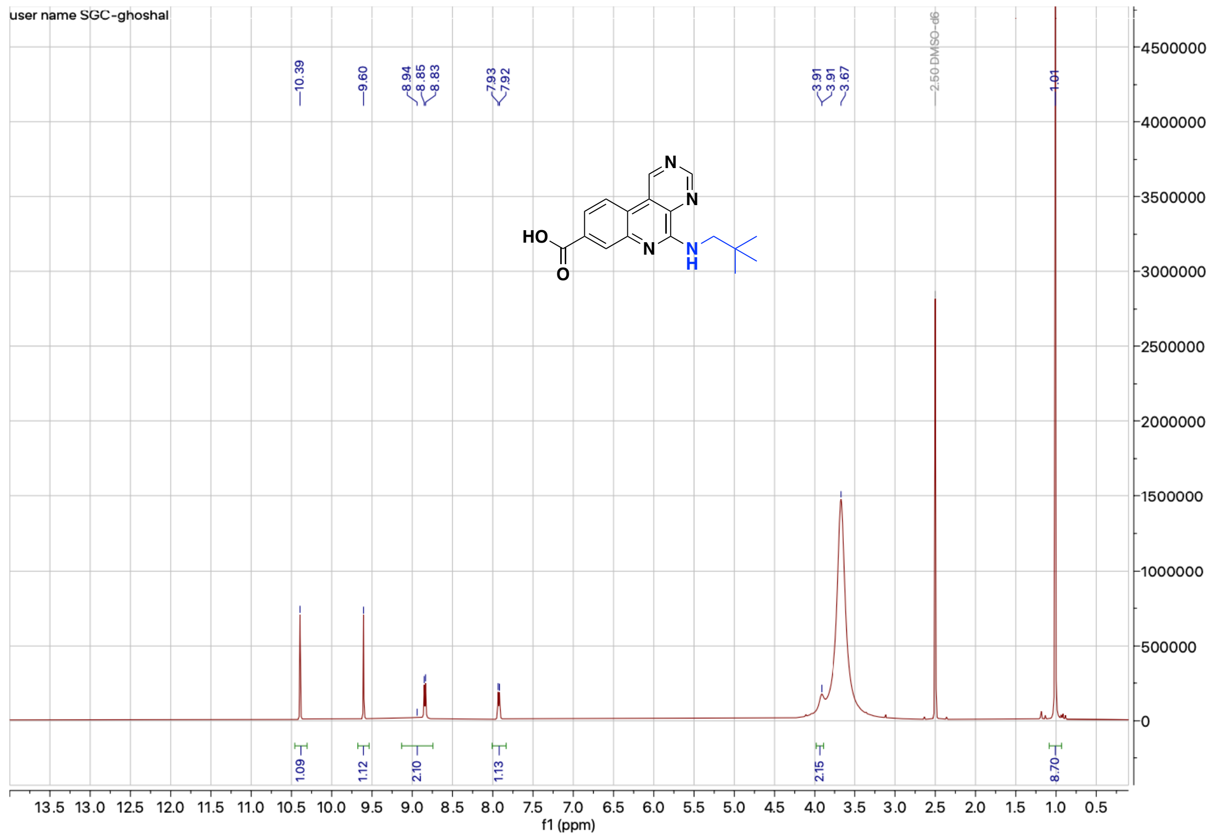


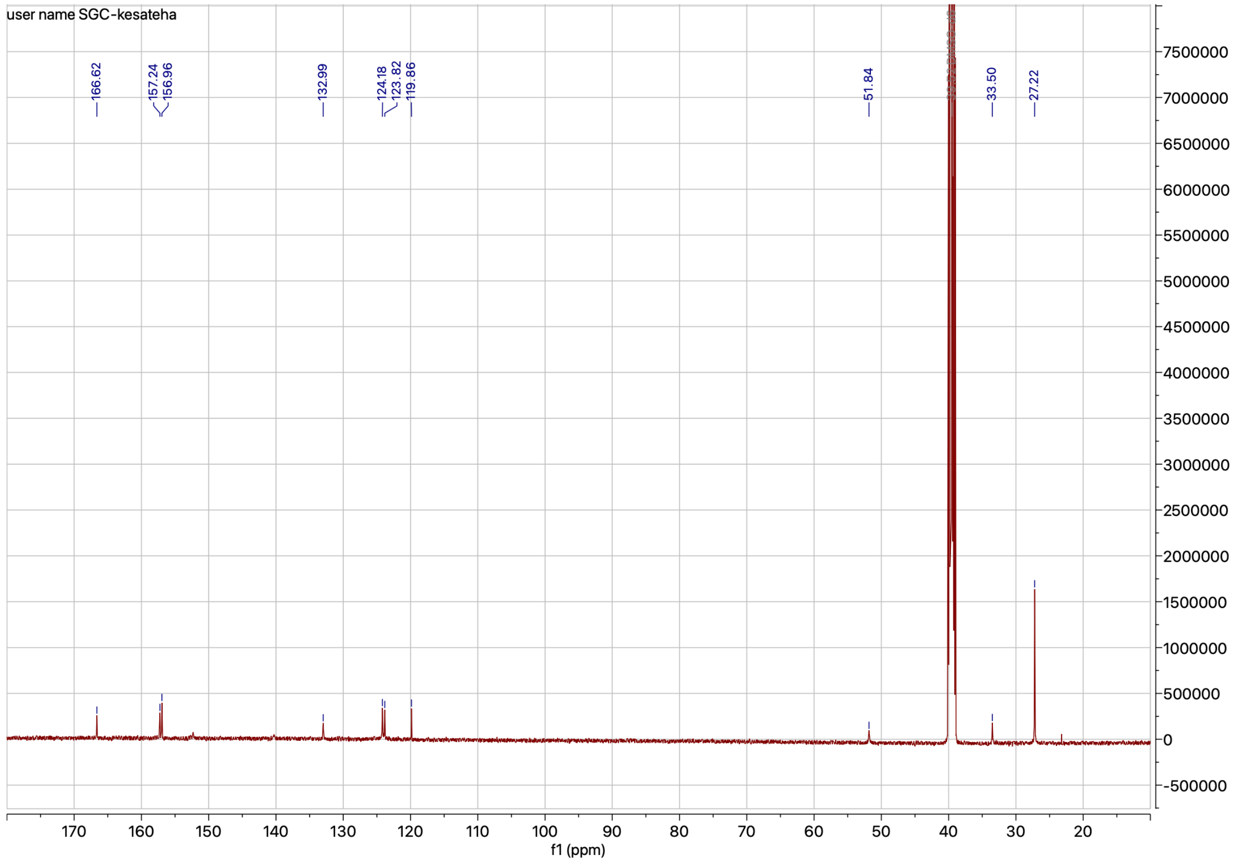


**Figure S24.** $^{1}$H and $^{13}$C NMR spectrum of compound **4v**

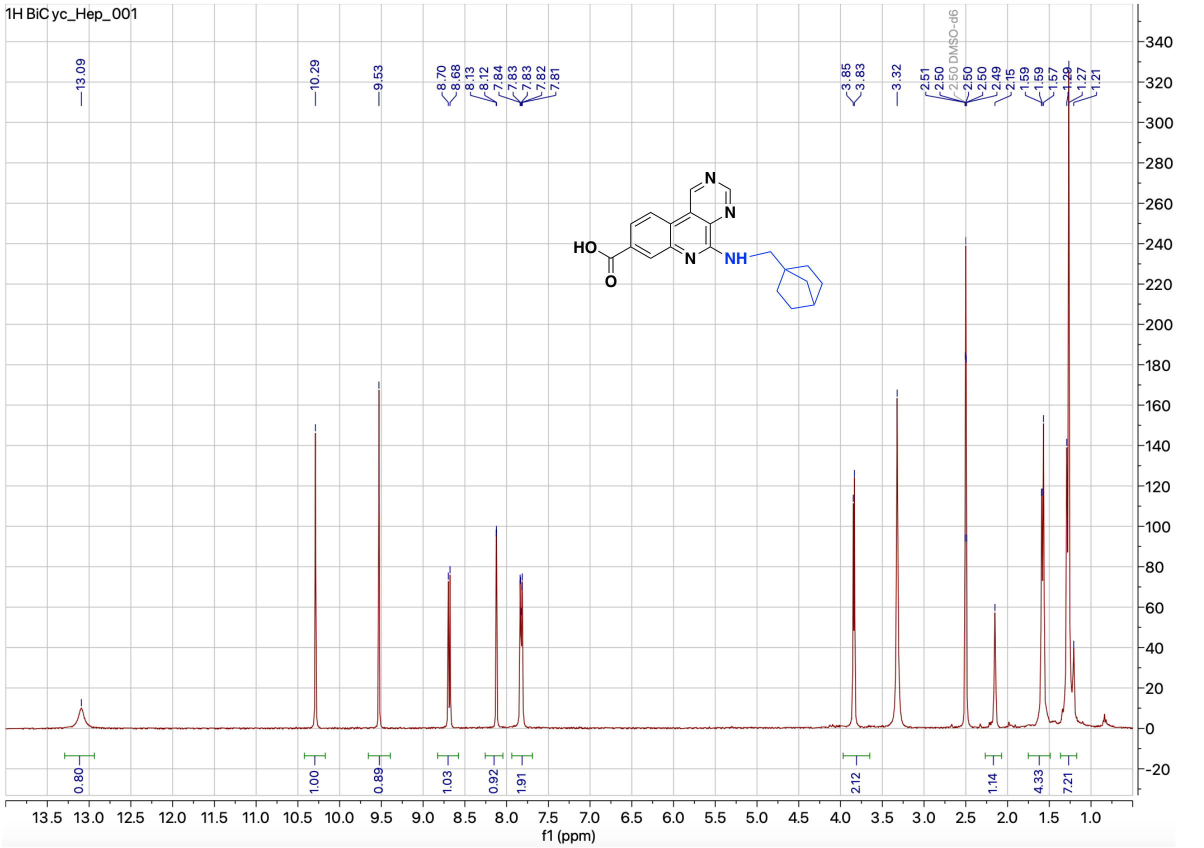

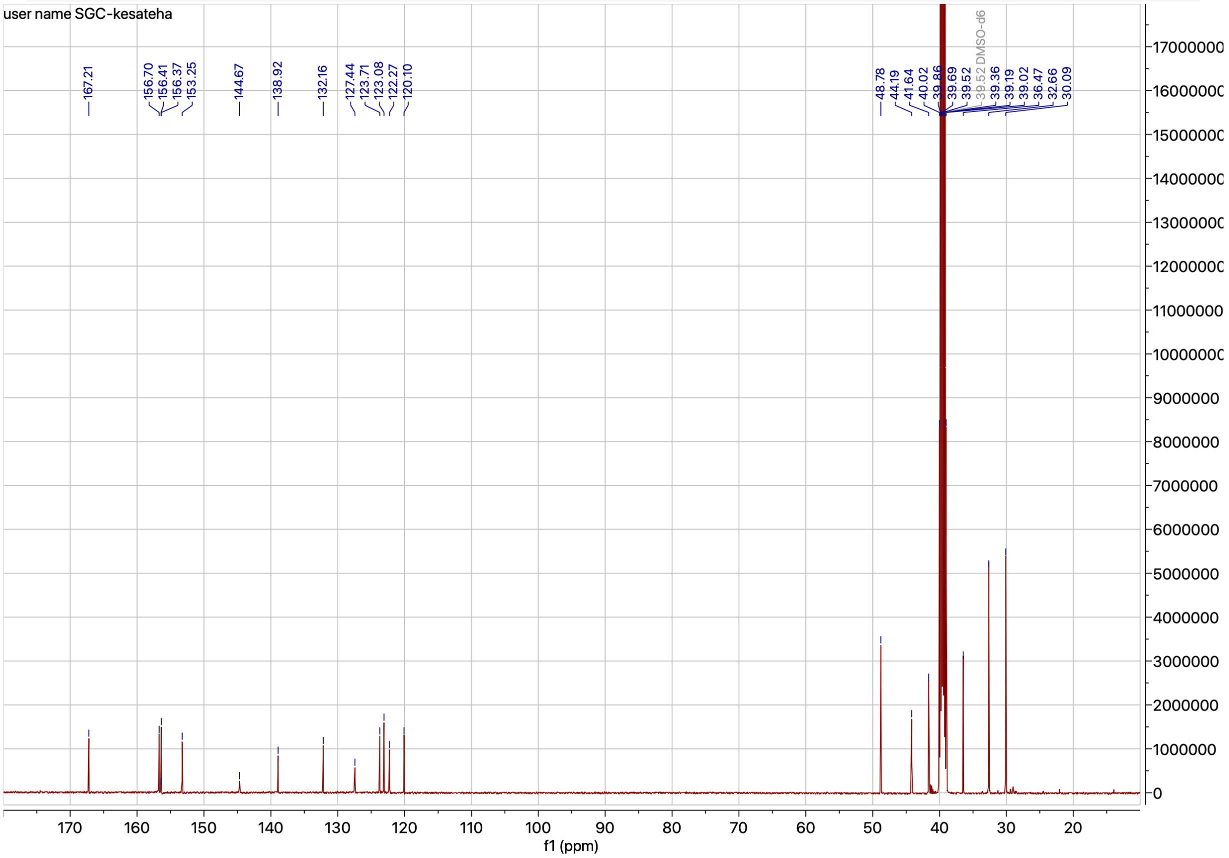

**Figure S25.** $^{1}$H and $^{13}$C NMR spectrum of compound **4w**

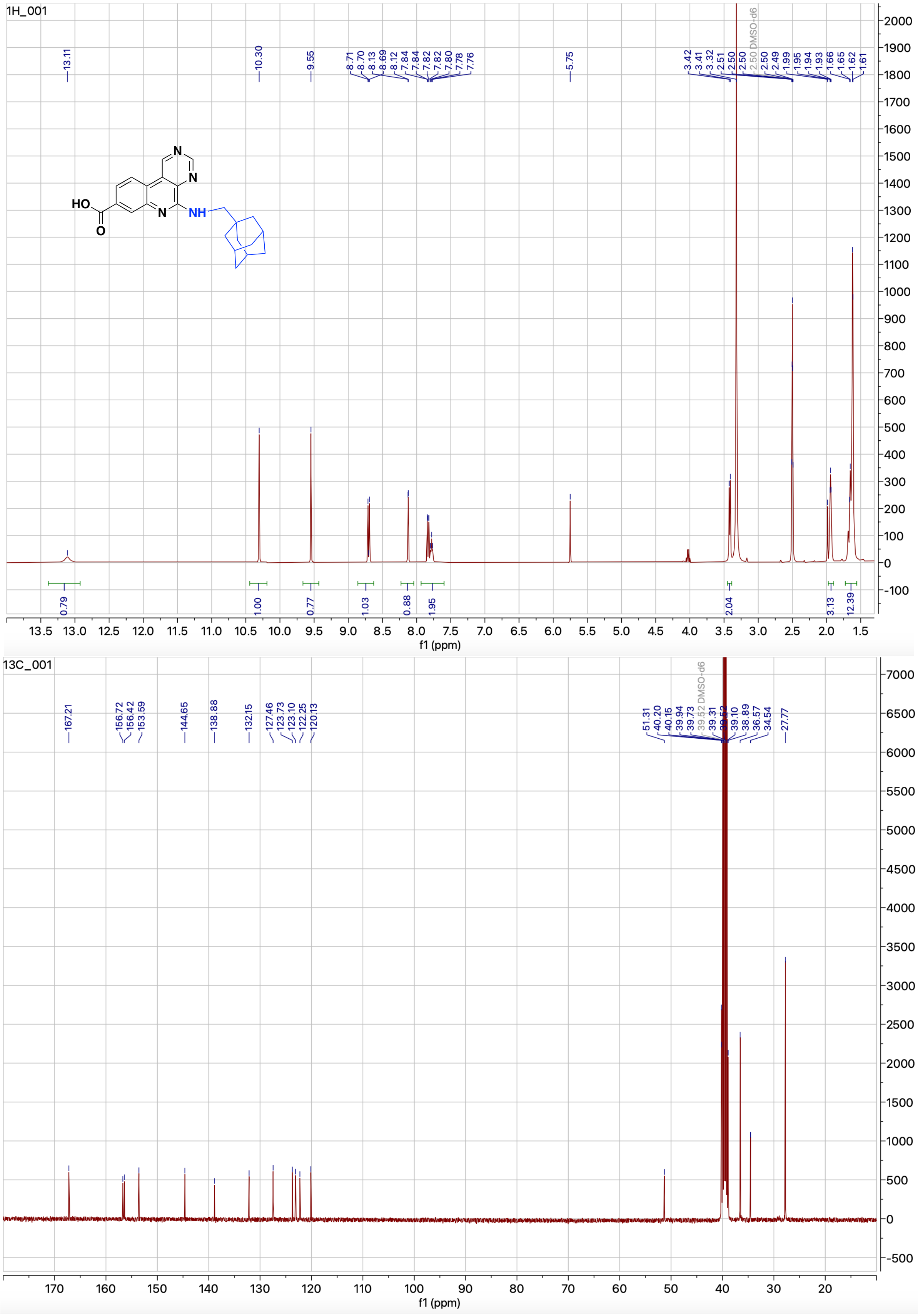


**Figure S26.** $^1$H and $^{13}$C NMR spectrum of compound **4x**

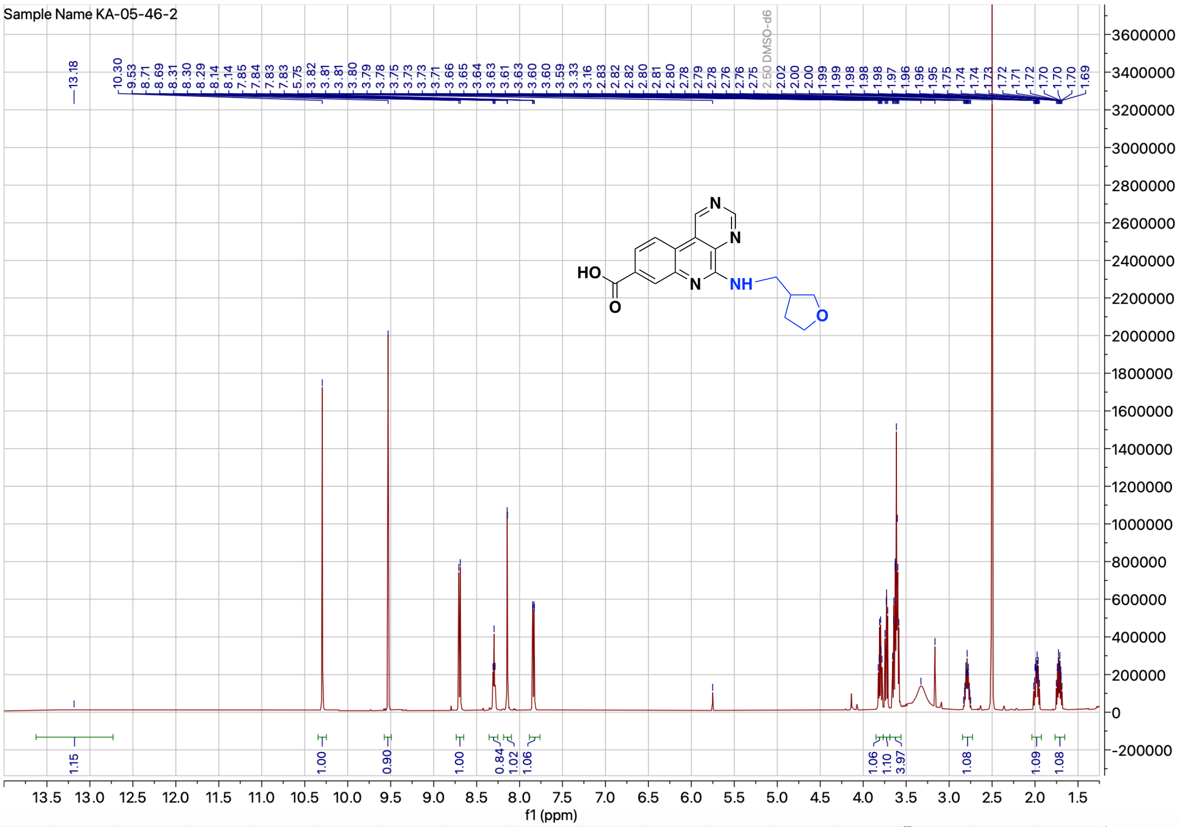


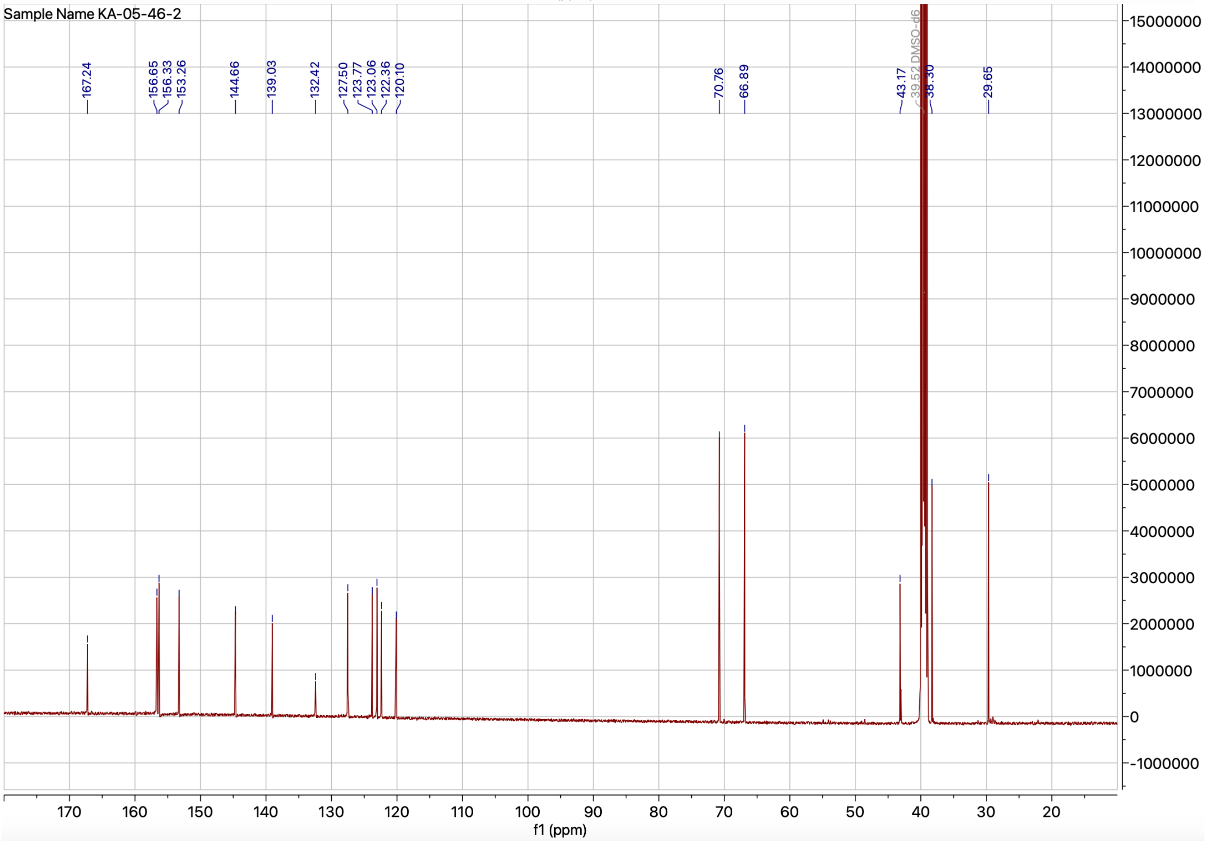


**Figure S27.** $^{1}H$ and $^{13}C$ NMR spectrum of compound **4y**

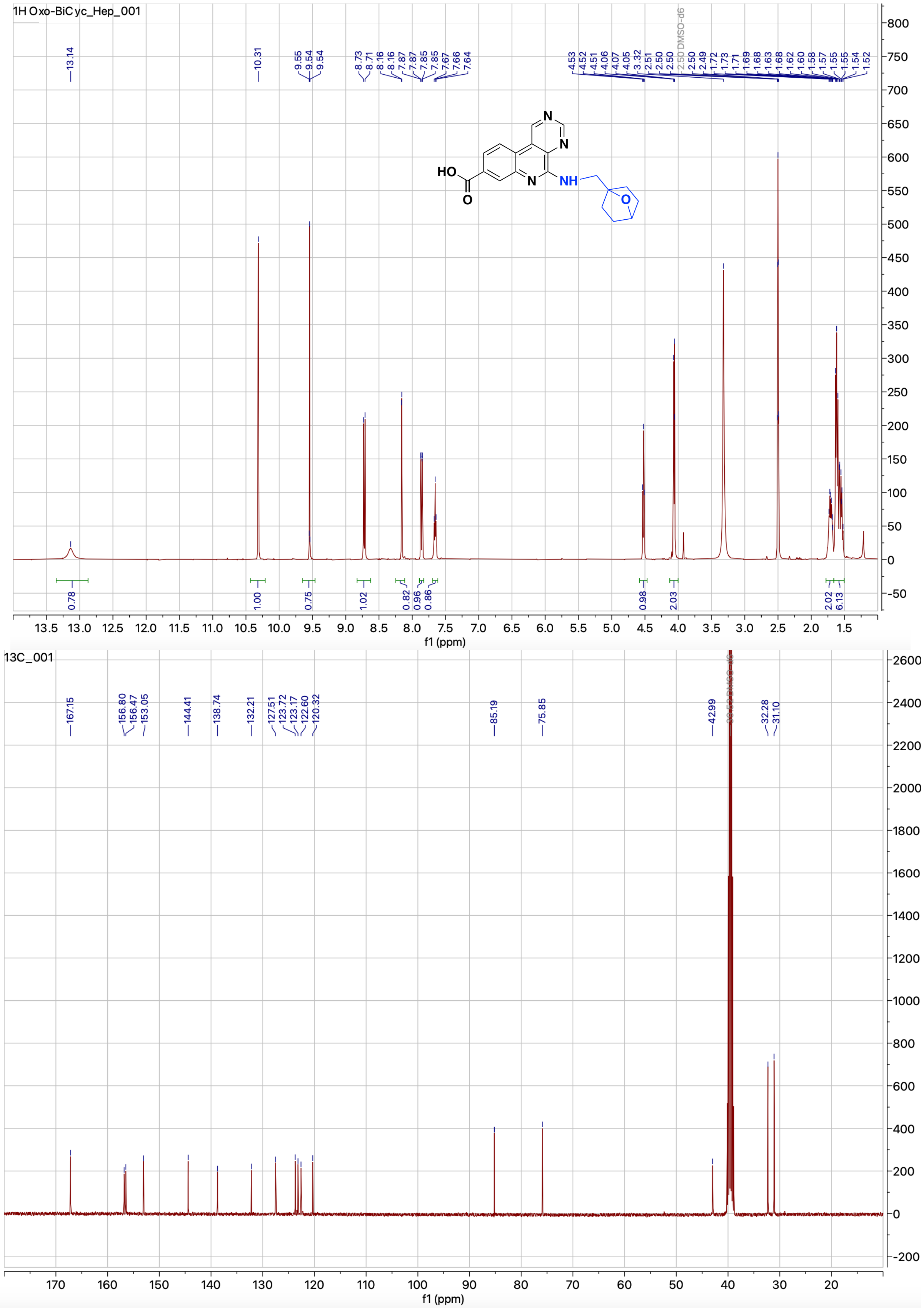


**Figure S28.** $^{1}$H and $^{13}$C NMR spectrum of compound **4z**

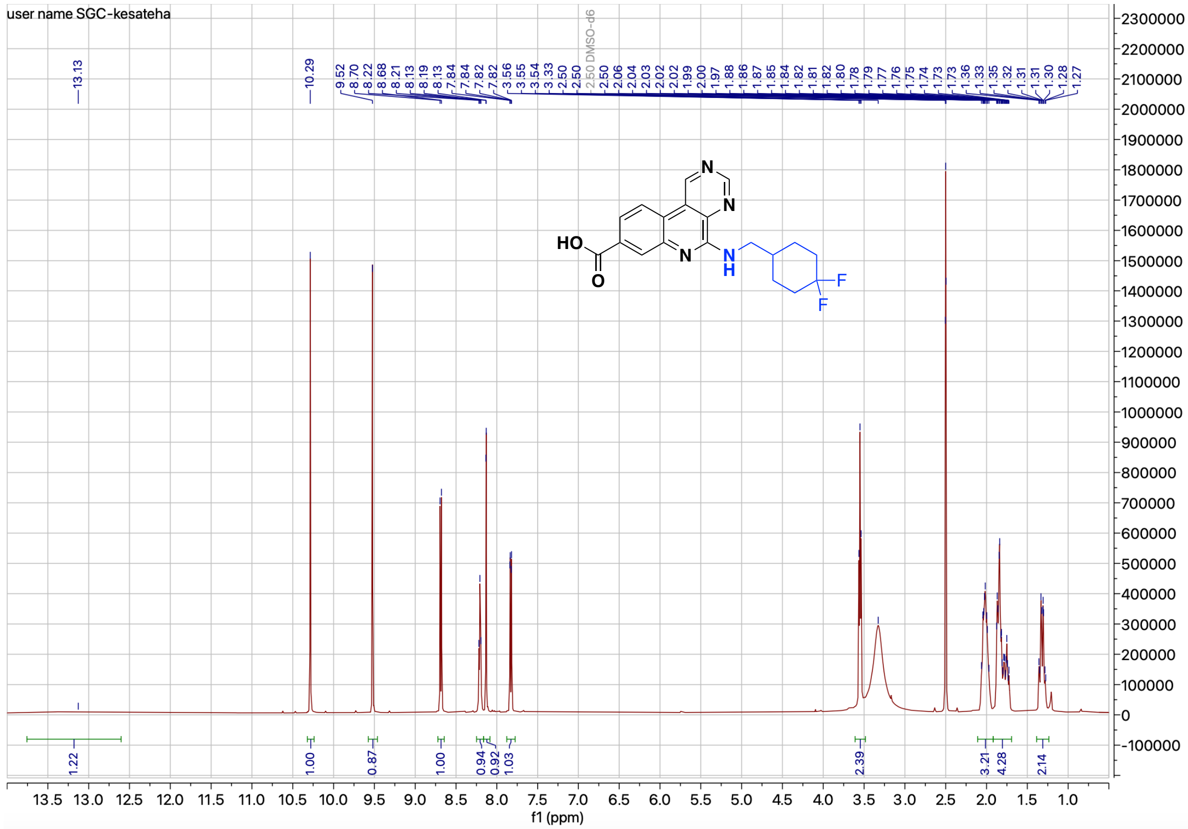


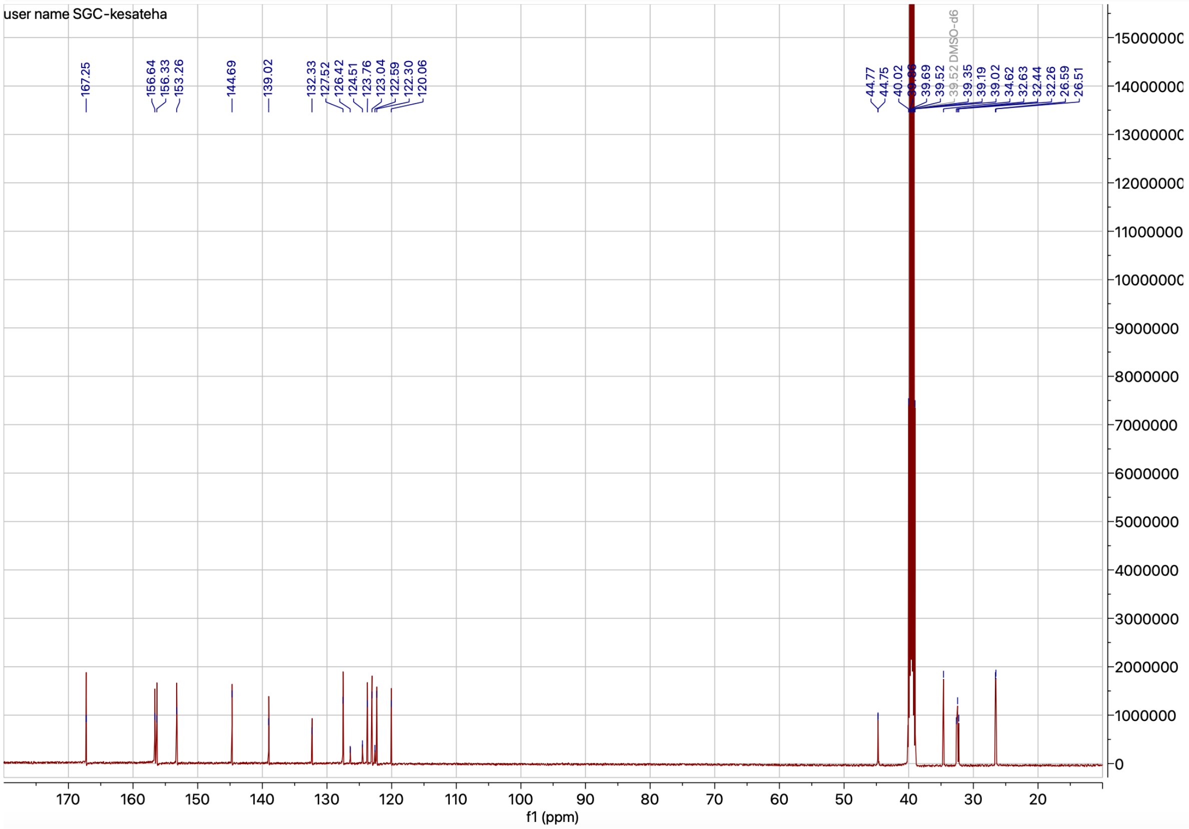


**Figure S29.** $^{1}$H and $^{13}$C NMR spectrum of compound **4aa**

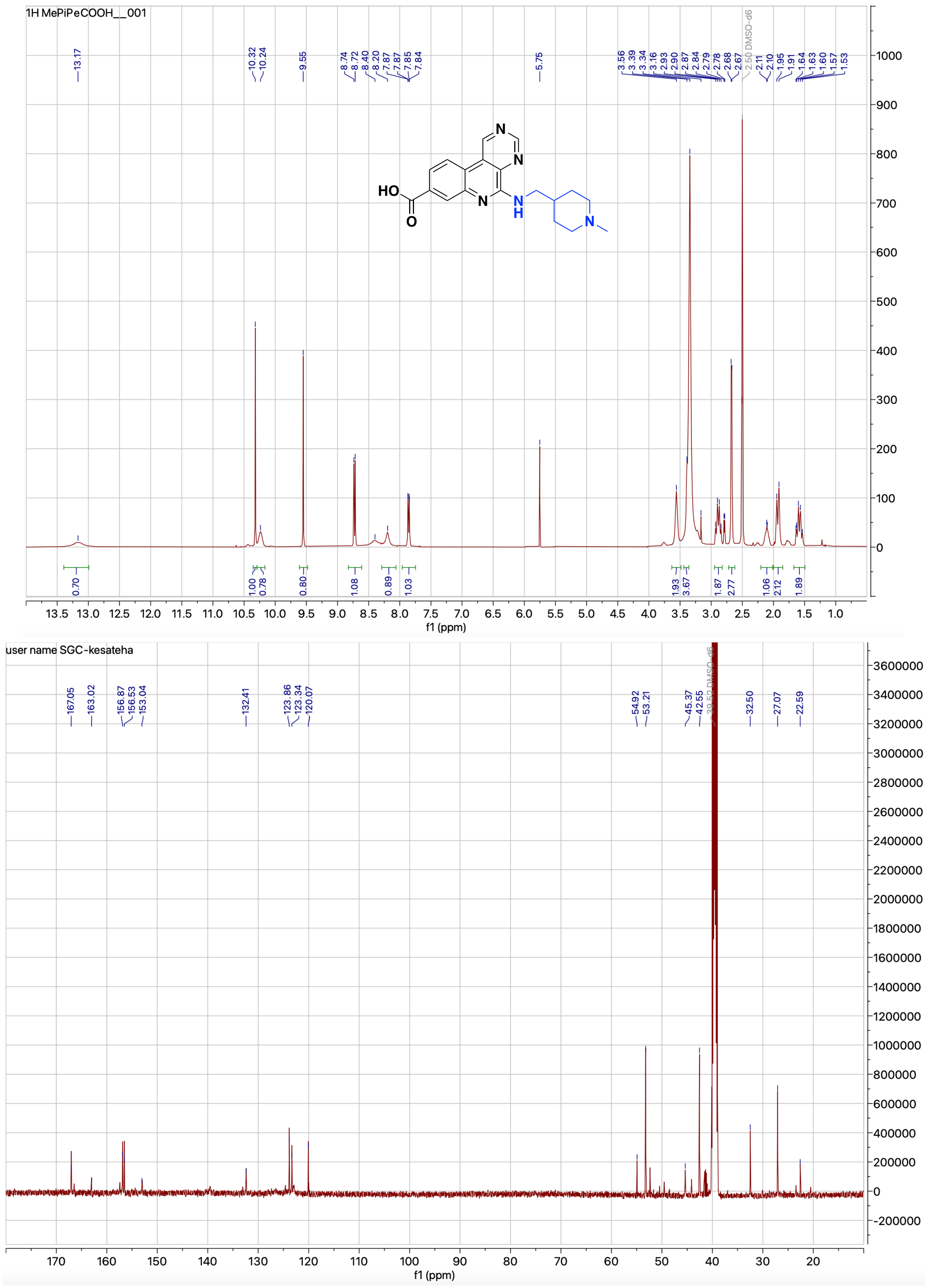


**Figure S30.** $^{1}$H and $^{13}$C NMR spectrum of compound **4ab**

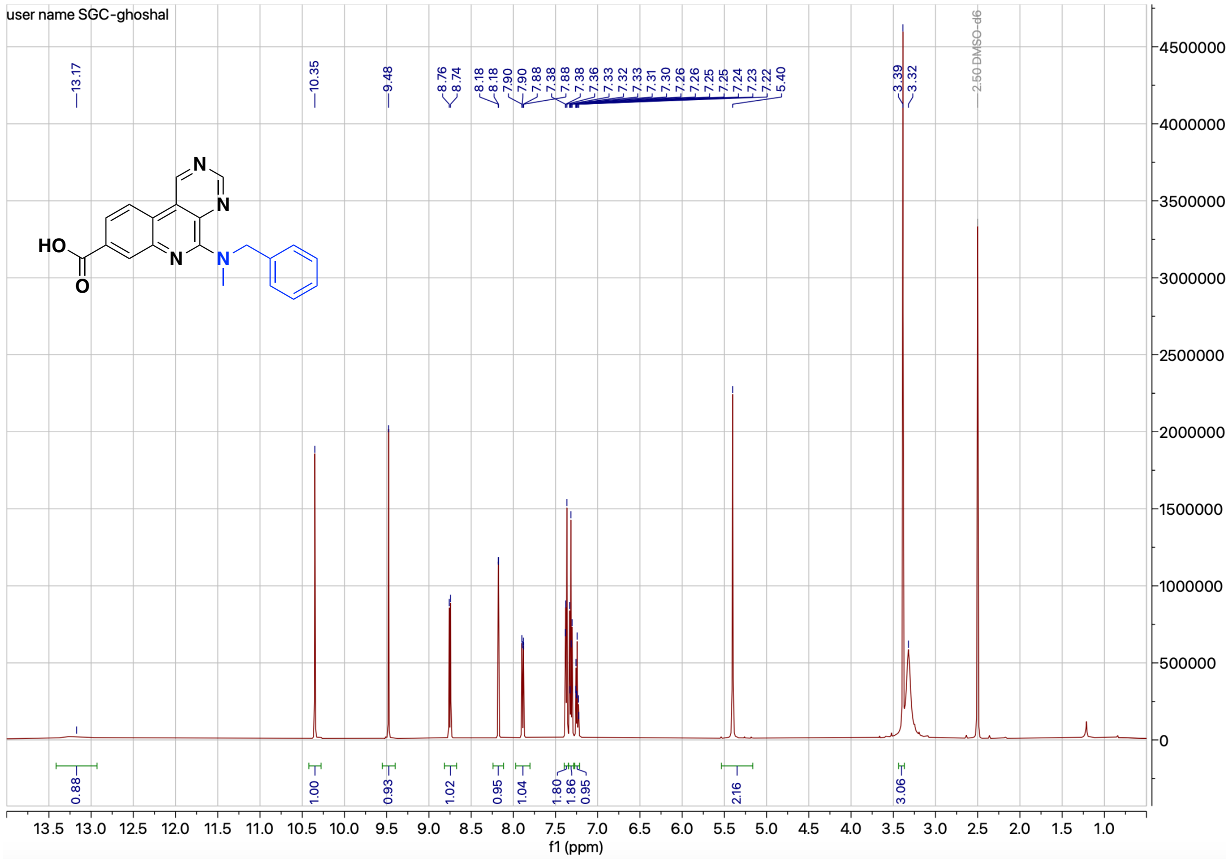

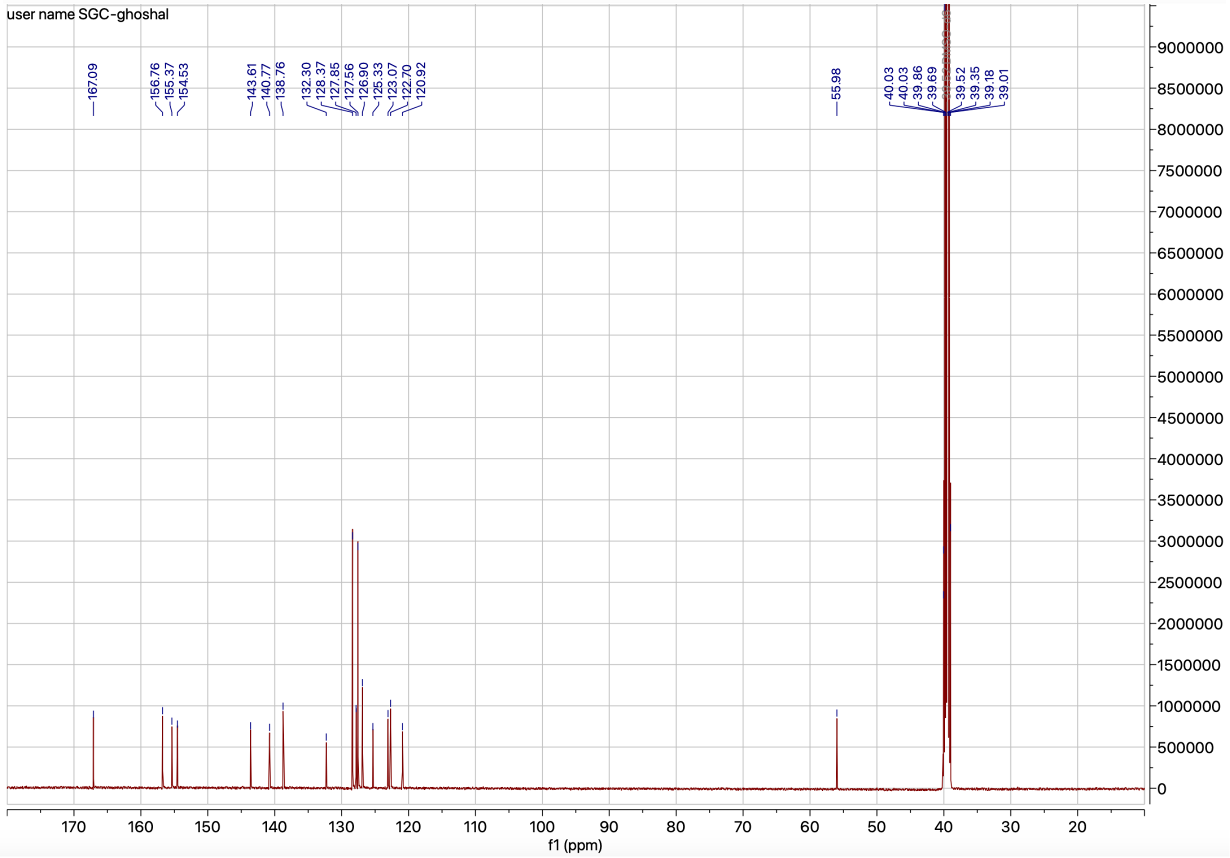

**Figure S31.** $^{1}$H and $^{13}$C NMR spectrum of compound **4ac**

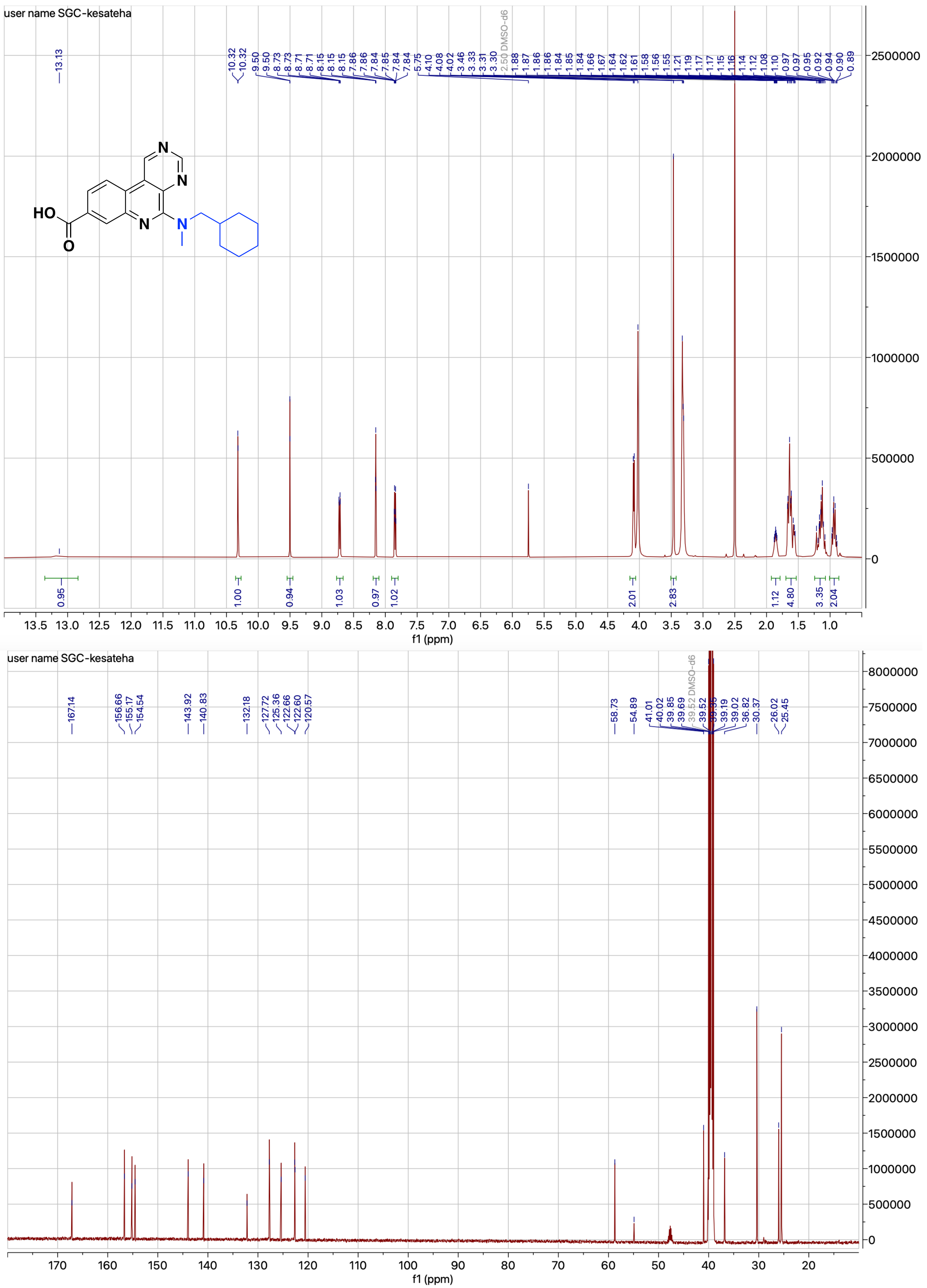


**Figure S32.** $^{1}$H and $^{13}$C NMR spectrum of compound **4ad**

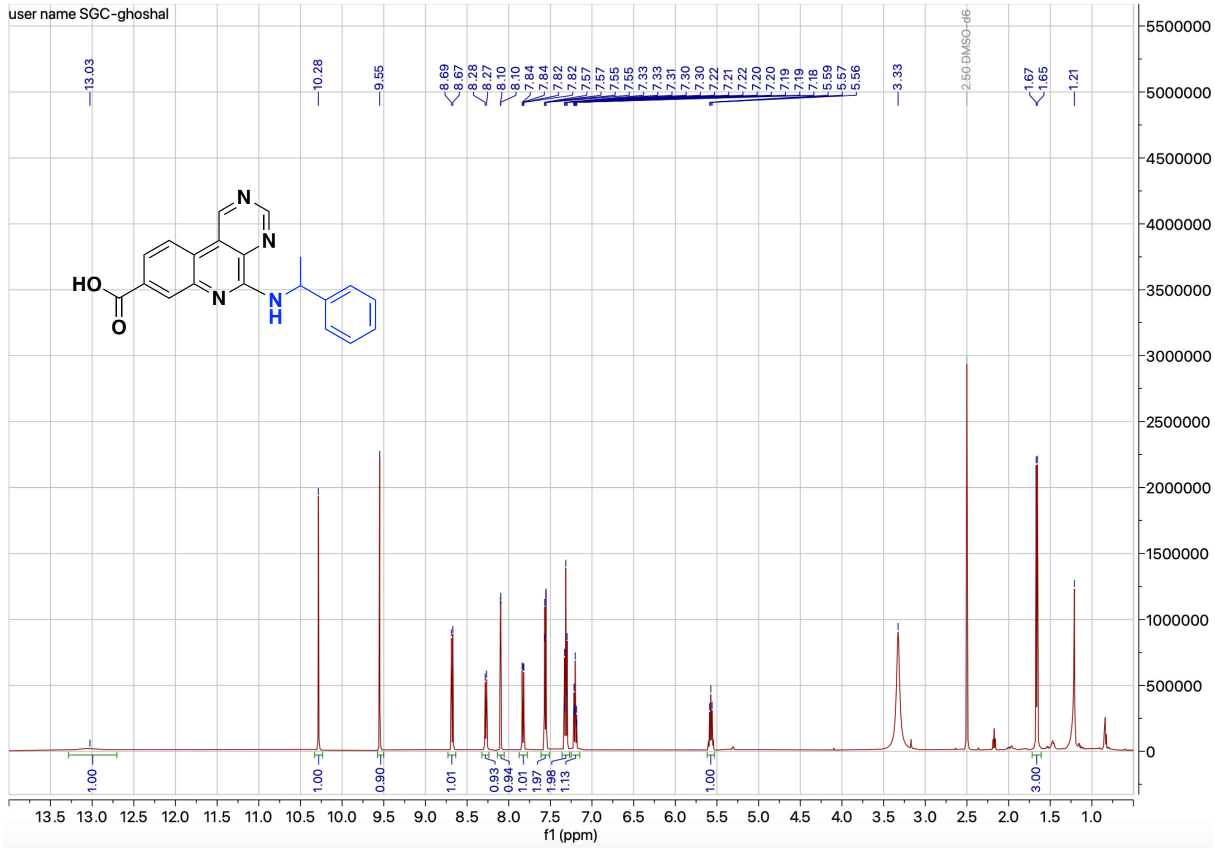


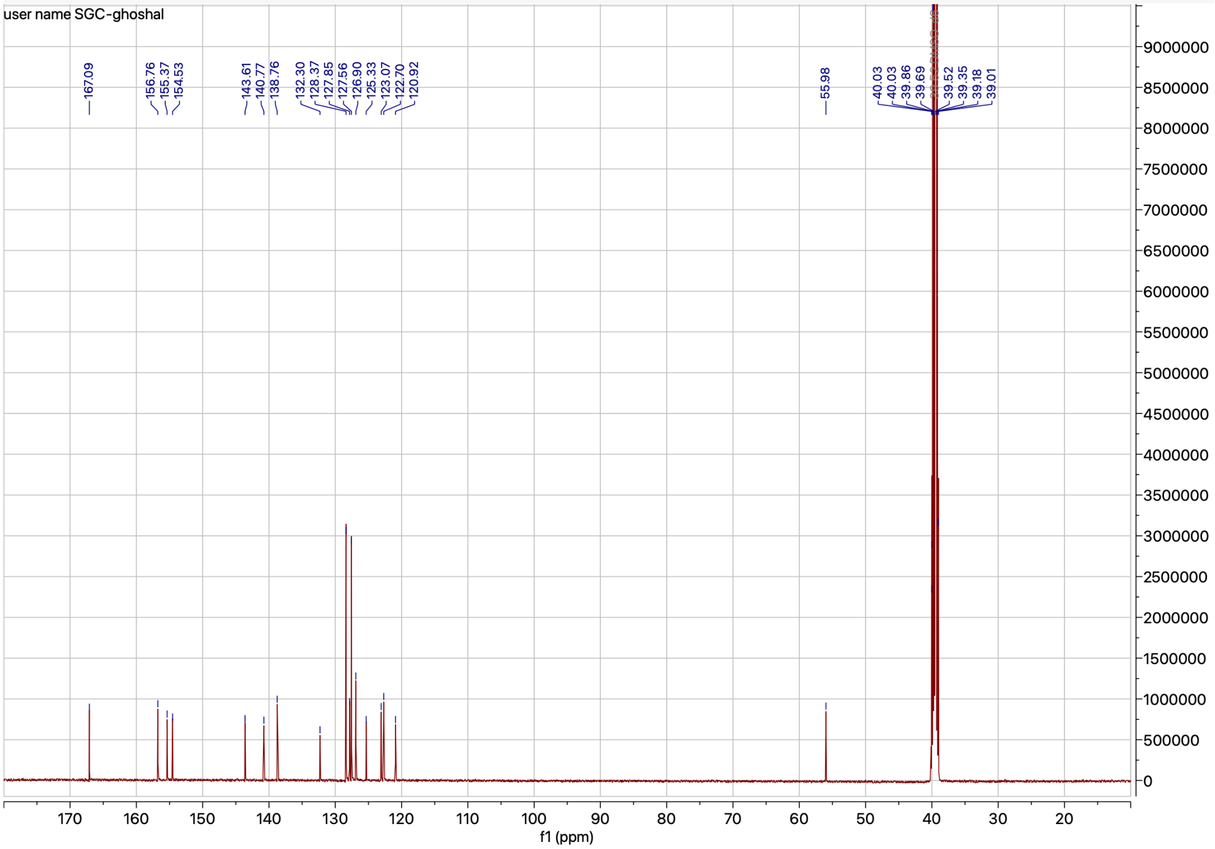


**Figure S33.** $^{1}$H and $^{13}$C NMR spectrum of compound **4ae**

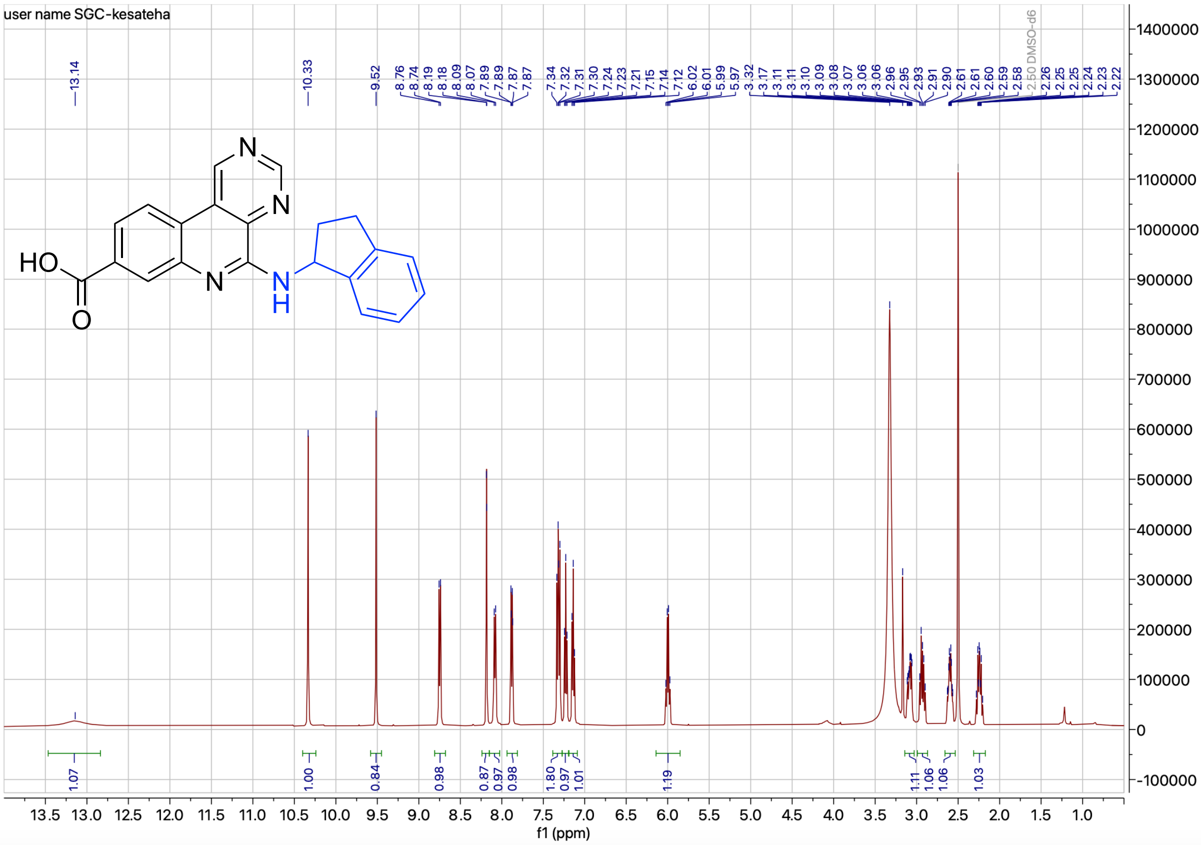


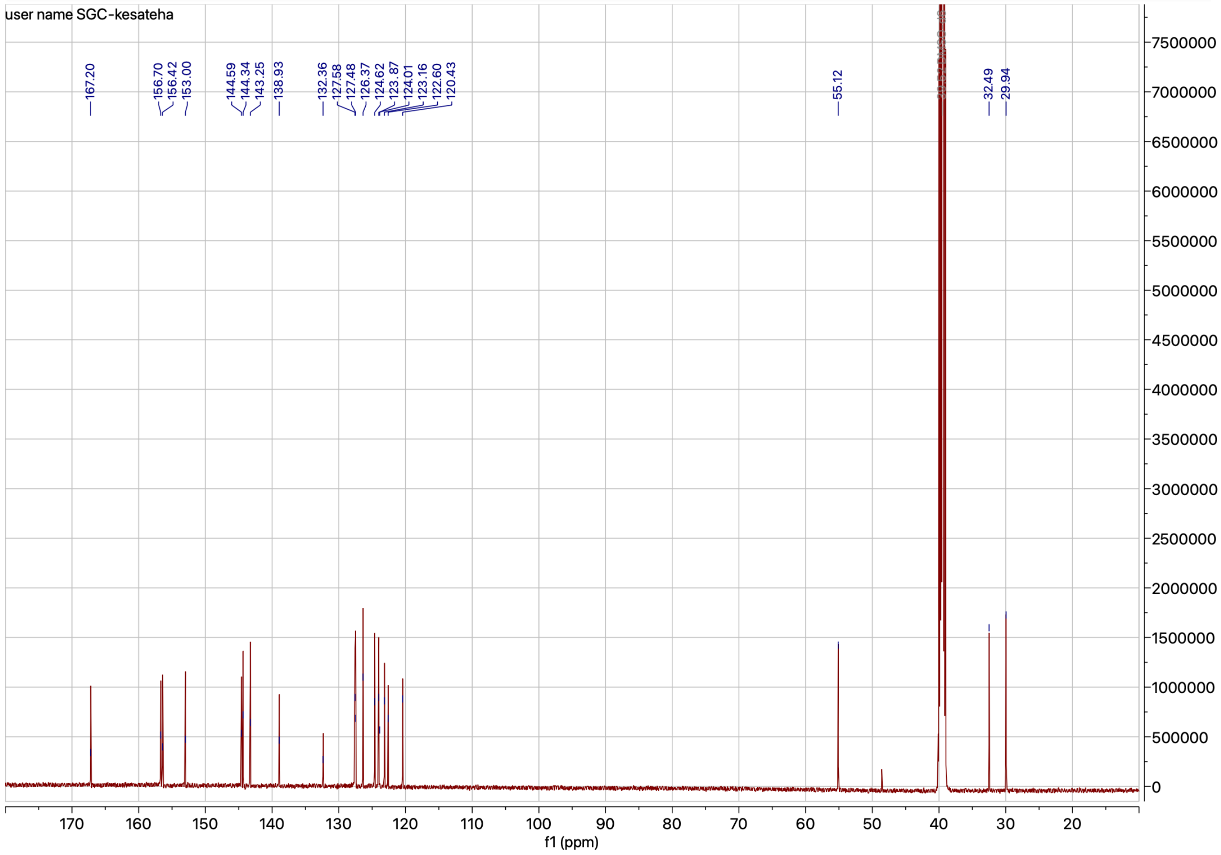


**Figure S34.** $^{1}$H and $^{13}$C NMR spectrum of compound **4af**

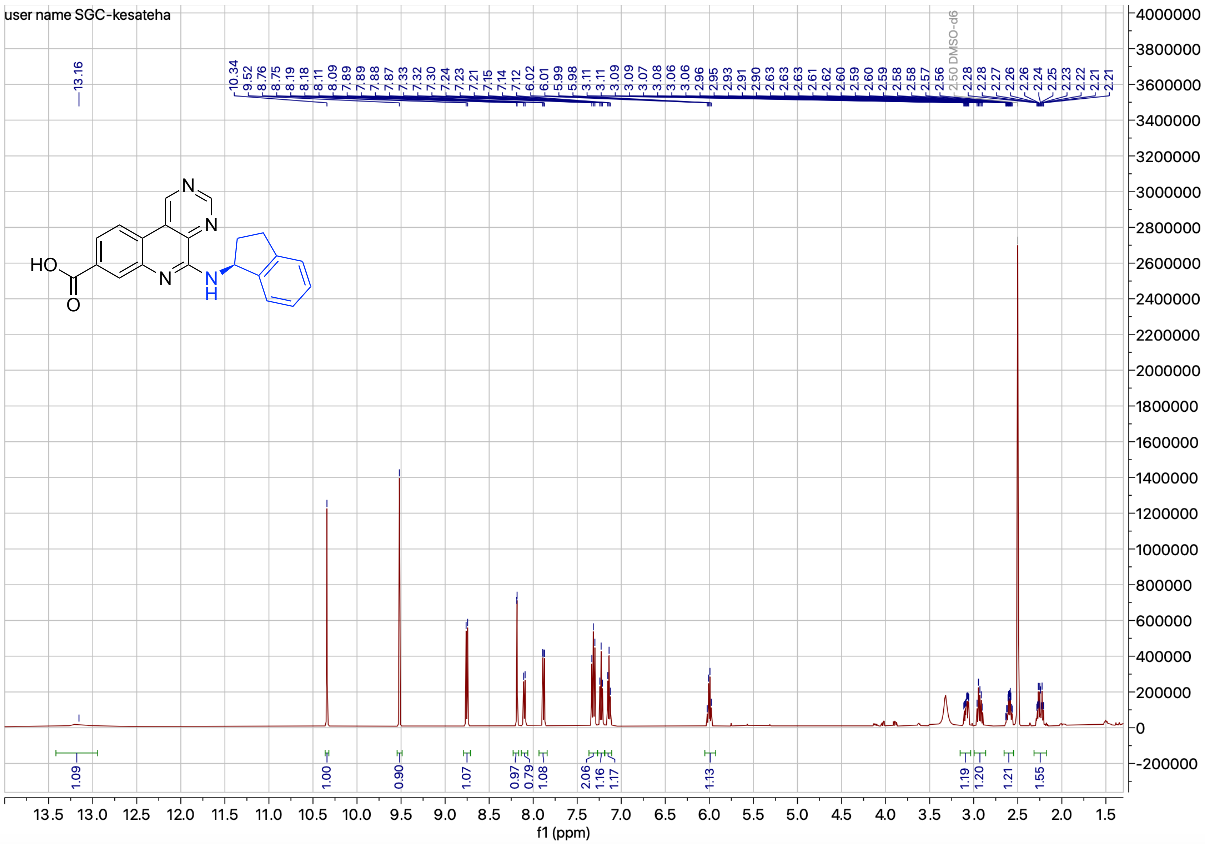

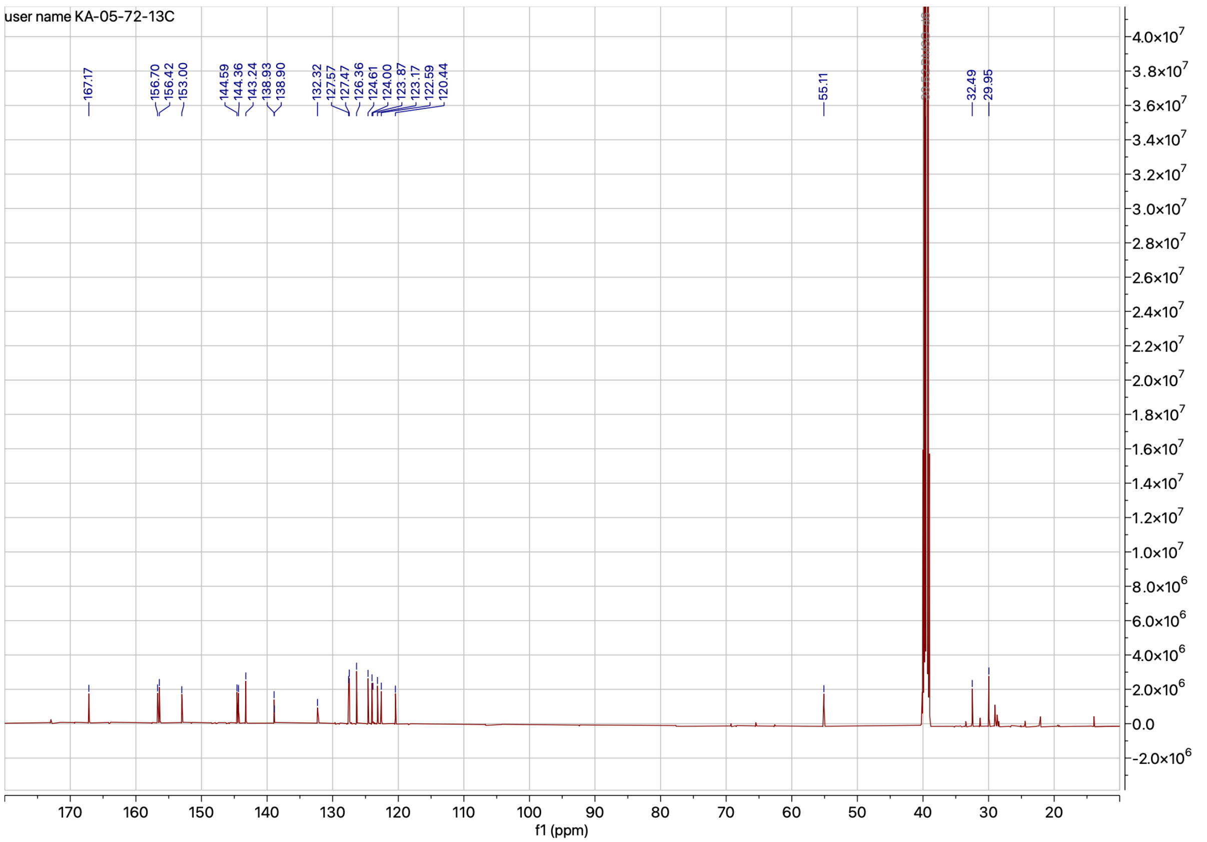

**Figure S35.** $^{1}H$ and $^{13}C$ NMR spectrum of compound **4ag**

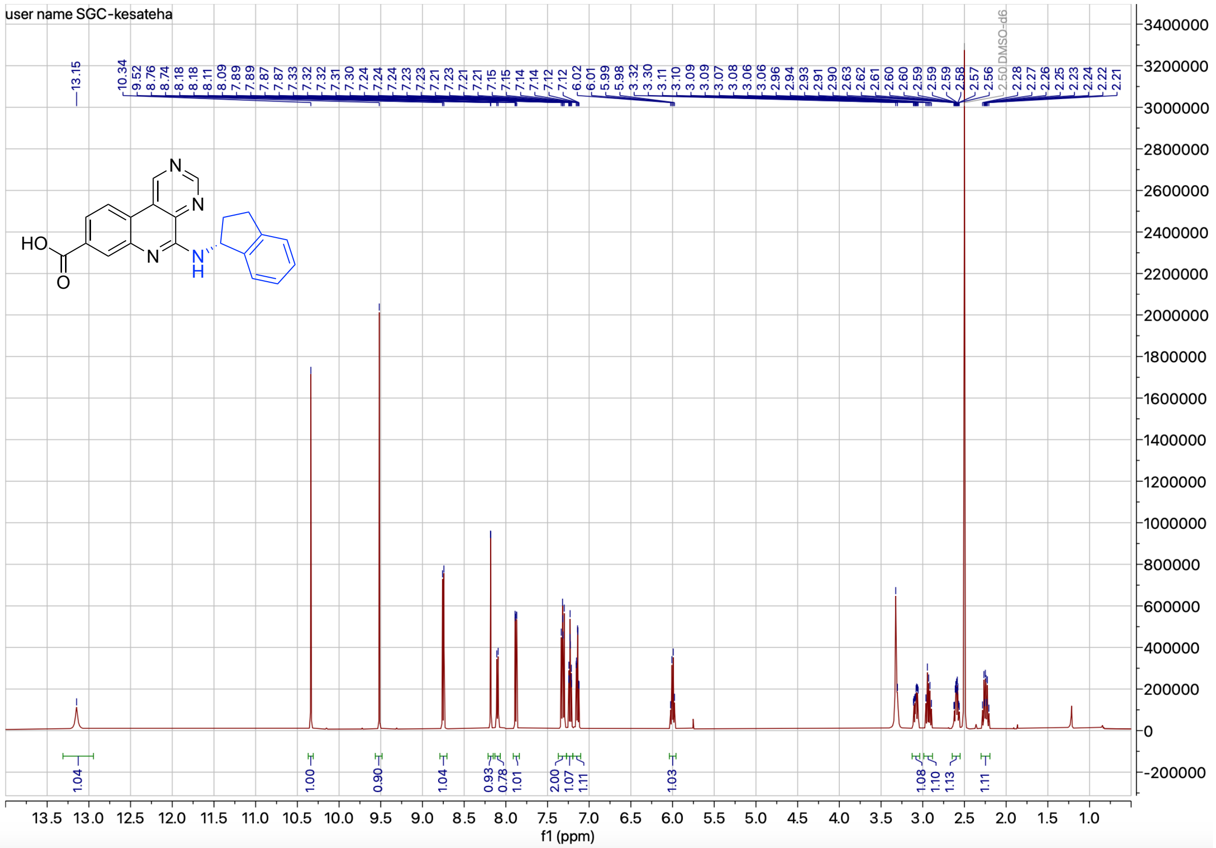

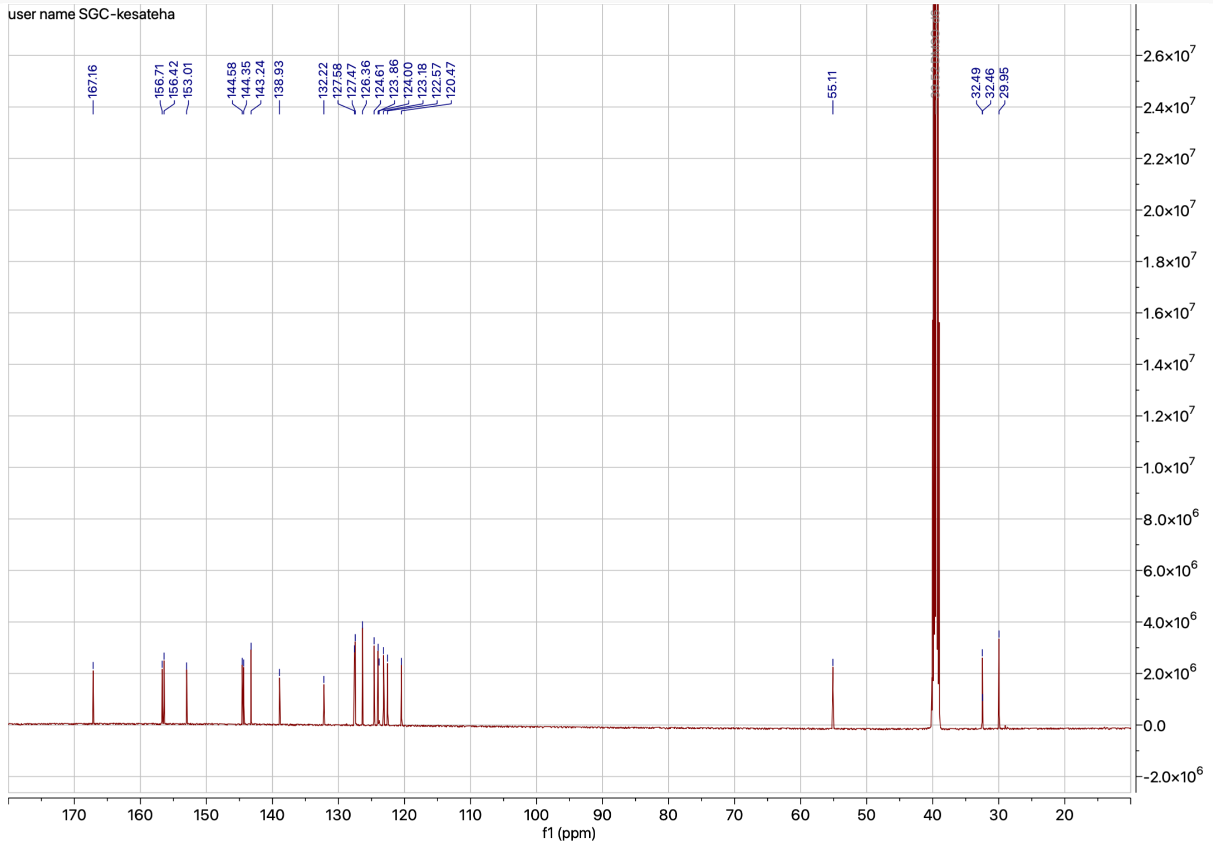

**Figure S36.** $^1$H and $^{13}$C NMR spectrum of compound **4ah**

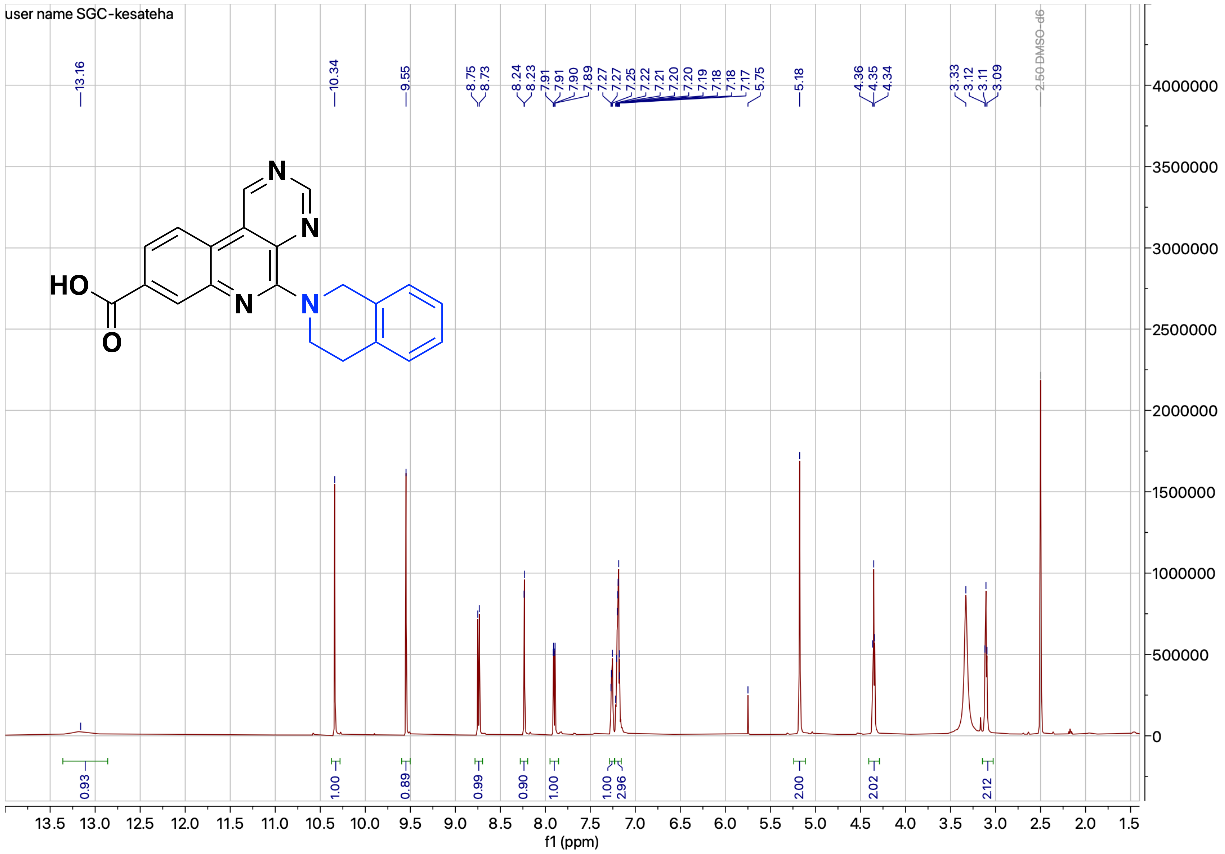

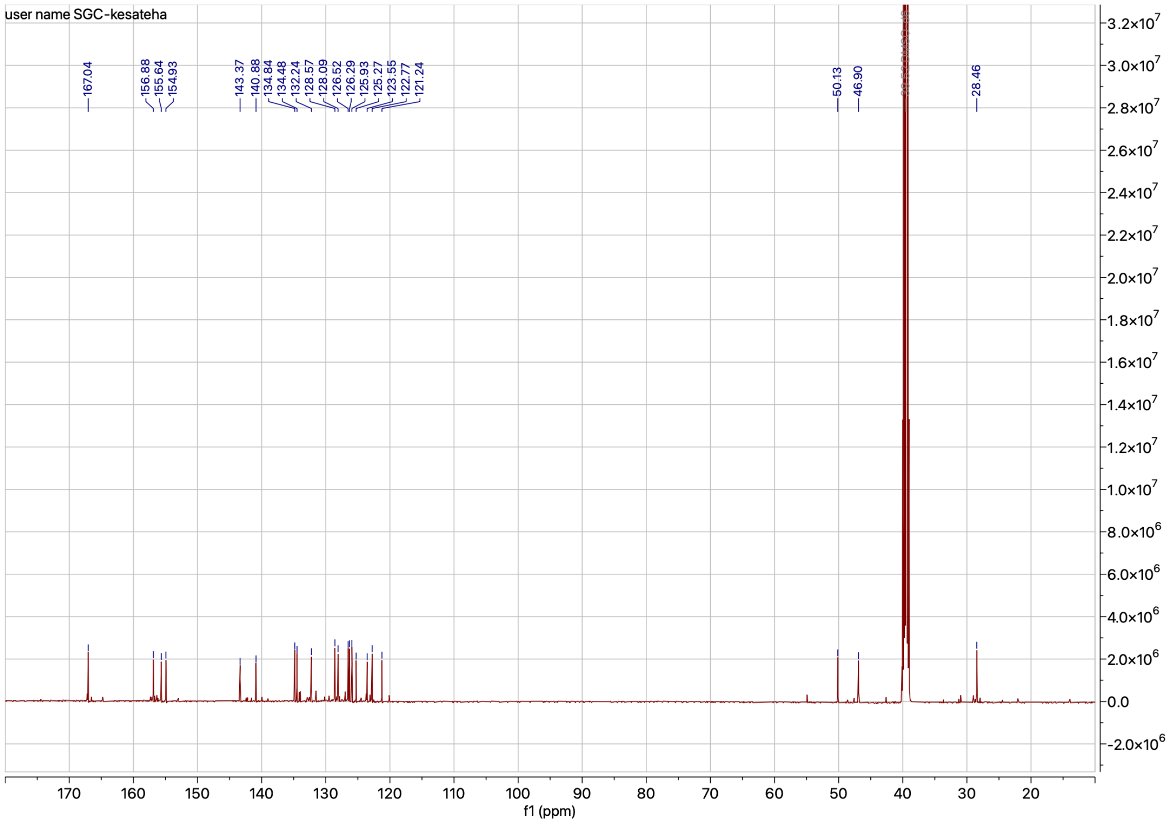

**Figure S37.** $^{1}$H and $^{13}$C NMR spectrum of compound **4ai**
